# Stochastic motion and transcriptional dynamics of pairs of distal DNA loci on a compacted chromosome

**DOI:** 10.1101/2023.01.18.524527

**Authors:** David B. Brückner, Hongtao Chen, Lev Barinov, Benjamin Zoller, Thomas Gregor

**Author notes:** These authors contributed equally (David B. Brückner, Hongtao Chen).

## Abstract

Chromosomes in the eukaryotic nucleus are highly compacted. However, for many functional processes, including transcription initiation, the 3D pair-wise motion of distal chromosomal elements, such as enhancers and promoters, is essential and necessitates dynamic fluidity. Therefore, the interplay of chromosome organization and dynamics is crucial for gene regulation. Here, we use a live imaging assay to simultaneously measure the positions of pairs of enhancers and promoters and their transcriptional output in the developing fly embryo while systematically varying the genomic separation between these two DNA loci. Our analysis reveals a combination of a compact globular organization and fast subdiffusive dynamics. These combined features cause an anomalous scaling of polymer relaxation times with genomic separation and lead to long-ranged correlations compared to existing polymer models. This scaling implies that encounter times of DNA loci are much less dependent on genomic separation than predicted by existing polymer models, with potentially significant consequences for eukaryotic gene expression.

---

Living systems are built based on information encoded in chromosomes confined in each cell’s nucleus. These meter-long DNA polymers must be highly compacted to fit into the micron-sized structure [1, 2]. At the same time, for cells to function, chromosome organization must allow the information content to be accessed and read out through transcription [3, 4]. Often transcription can only occur through the spatial interaction of DNA loci, such as enhancers and promoters. They find each other dynamically and remain in physical proximity [5–8]. While the distances over which many enhancers function in higher eukaryotes can be up to mega-base pairs in genomic separation [9–12], it is unknown how these elements come into proximity, what their typical distance is in 3D space, and how they explore this space dynamically in the process. Specifically, it remains unclear how the real-time physical motion of such coupled pairs of DNA loci determines transcriptional encounters and how this depends on their genomic separation.

Over the past decade, the advent of chromosome capture and imaging methods [13] has given key insights into the 3D spatial organization of chromosomes, with the discovery of structural features such as topologically associating domains (TADs) [14–17], phase-separated nuclear condensates [18–20], and larger-scale compartments [21, 22]. These organizing structures have key implications for transcriptional regulation [23]. However, these structures are not static, but have been revealed to be heterogeneous across cells [24, 25] and dynamic and short-lived in time [26, 27]. The role of the real-time dynamics of pairs of loci is only beginning to be understood and remains elusive for focal contacts which are key to establishing enhancer–promoter interactions in many systems [28].

Similarly, from a polymer physics perspective, there is a gap in our understanding of static and dynamic properties of chromosomes. At large scales, across tens to hundreds of TADs, chromosome organization has been suggested to be highly compacted in a crumpled chain configuration (also referred to as fractal globule), a long-lived polymer state with fractal dimension three [22, 29–32]. Yet, the real-time dynamics of DNA loci revealed by live-imaging experiments exhibit subdiffusion with exponents close to the predictions of the simple Rouse polymer model [33], which predicts a loosely packed ideal chain polymer configuration with fractal dimension two that contrasts the compacted architecture of the crumpled chain model [26, 27, 34–36]. A promising technique to address this gap are scaling approaches that combine fractal organization and subdiffusive dynamics [37–39], but these have never been tested experimentally.

Thus far, experimental data sets have given insight into either static organization [14–17, 22, 32], dynamic properties of chromosomes [26, 27, 34, 35, 40], or transcription [8, 36, 41–43], but rarely all at the same time. For instance, previous live measurements of locus pairs occurred at fixed genomic separation in transcriptionally silent loci [26, 27]. To investigate how 3D spatial organization and dynamic locus motion control the encounter times of functional DNA loci and thus transcriptional activation, we require an approach to simultaneously monitor the movement of DNA locus pairs and transcription across a series of genomic separations *in vivo*.

Here, we address this problem by live imaging the joint dynamics of two cis-regulatory DNA elements, an enhancer, and a promoter, while simultaneously monitoring the transcriptional output resulting from their functional dynamic encounters in developing fly embryos. We systematically vary the genomic separation between these loci spanning many TADs. Stochastic real-time trajectories of the 3D motion of the two loci show a dynamic search process, with physical proximity required for successful transcription and a power-law scaling of transcription probability with genomic separation. While typical 3D distances between the locus pair follow a compact packing consistent with the crumpled chain model, the dynamic properties exhibit fast diffusion, albeit with a diffusion coefficient that increases with genomic separation. These features give rise to an anomalous scaling of polymer relaxation times and long-range correlations in the relative motion of the two loci. This suggests that the enhancer–promoter search process is much less dependent on genomic separation than expected based on existing polymer models.

## Live imaging of chromosome dynamics and transcription

To simultaneously monitor the coupled motion of enhancer–promoter pairs and transcription across multiple genomic separations, we generated fly lines, in which a reporter gene is introduced at various genomic locations from the well-studied *Drosophila even-skipped* (*eve*) locus (Fig. 1b). The locations of both the endogenous *eve* enhancers and the promoter of the reporter gene, as well as the transcriptional activity of the reporter gene are measured together using a three-color imaging system (Fig. 1a, Supplementary Section S1.2) [8]. To facilitate transcription, the reporter cassette contains the insulator element *homie*, which allows stable loop formation with the endogenous *homie* element in the *eve* locus (Fig. 1b).

**Figure 1.**
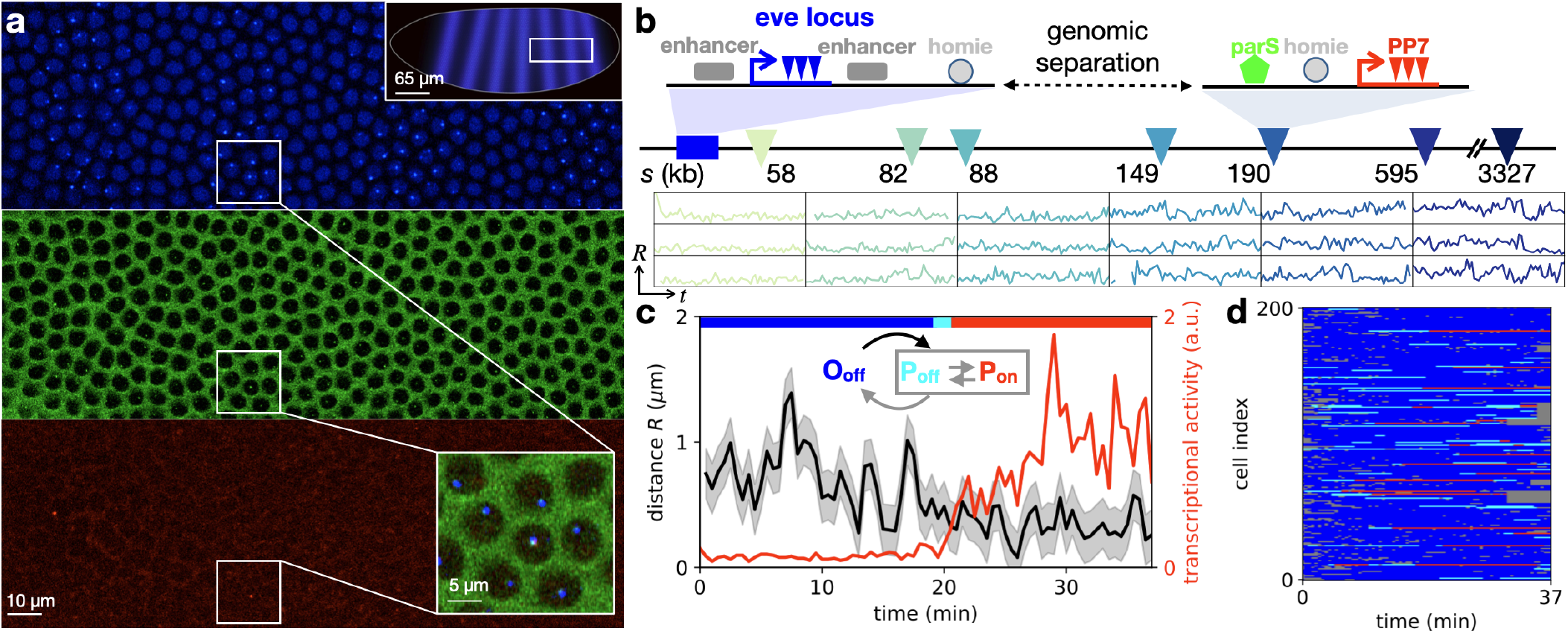
Simultaneous tracking of DNA loci and transcriptional activity in living embryos. **a**, Typical surface view of a representative fly embryo, displaying fluorescent foci for MS2, parS, and PP7 in the corresponding blue (top), green (center), and red (bottom) channels. Top inset shows schematic with image location in the embryo; bottom inset shows a close-up. **b**, Top: schematic of the gene cassettes used for three-color imaging. The endogenous *eve* locus (left) is tagged with MS2 stem-loops that are labeled via blue fluorescence. A reporter with an eve promoter driving PP7 transcription (labeled via red fluorescence) is integrated at a genomic separation *s* from the eve locus on the 2^nd^ chromosome in the *Drosophila* genome. It includes a homie insulator sequence allowing loop formation through homie–homie pairing, and a parS sequence that is permanently labeled with green fluorescence. Seven such constructs were generated with varying genomic separation *s* (triangles). Bottom: sample inter-locus distance trajectories *R*(*t*) for six genomic separations, with standardized y-axis limits (0, 2 μm) and x-axis limits (0, 30 min), obtained following nucleus and locus segmentation, tracking, chromatic aberration, and motion correction (Supplementary Section 1.1). The sampling time interval is 28 *s*. **c**, Trajectories of inter-locus distance *R* and transcriptional activity, with inferred topological states shown by the colored top bar (blue: O_off_, cyan: P_off_, red: P_on_; Supplementary Section 2). Inset: Schematic of the three topological states. **d**, 200 examples of state trajectories sampled from a total set of *N* = 579 trajectories acquired in *n* = 30 embryos (genomic separation *s* = 149 kb). Colors correspond to the legend of panel c; grey parts of the trajectories correspond to time points where the loci could not be tracked.

We build seven of such reporter constructs, with genomic separations *s* varying over close to two orders of magnitude from 58kb to 3.3Mb, comparable to the distances over which many enhancers function in higher eukaryotes (Supplementary Section S1.1) [9–12]. Importantly, these genomic length-scales span across multiple TADs in the *Drosophila* genome, with typical median TAD sizes of 90 kb [45] (here 18 kb for the *eve* locus).

Imaging took place for 30 min during the second half of nuclear cycle 14 (NC14) of embryo development (Fig. 1c), well after the completion of DNA replication. Sister chromatids are tightly coupled together at intervals < 10 kb [46]. Therefore, our two tagged DNA loci are connected by a single chromatin polymer composed of two coupled chromatids that are not resolved by our microscopy.

## Inter-locus distance scaling suggests crumpled chain organization

As demonstrated previously [8], this system exhibits three topological states (Fig. 1c): an open configuration O_off_ where the *homie* elements are not bound to each other, and two paired configurations P_off_ and P_on_, where a loop is formed with either inactive or active transcription, respectively. To determine the instantaneous topological and transcriptional state of the system, we adopt an inference approach using a Hidden Markov Model based on the time series of interlocus distance and transcriptional activity (Supplementary Section S2). We assign one of these states to each measured configuration, including the hidden P_off_ state (Fig. 1d).

A key question is how the inter-locus distances *R* in the open configuration O_off_ vary with the linear genomic separation *s*. These distances exhibit broad distributions, which shift systematically with larger separation (Fig. 2a). From a polymer physics perspective, the mean distance 〈*R*〉 is expected to scale as *s*^1/*d*^, where *d* is the fractal dimension: while an ideal chain polymer has fractal dimension *d* = 2, the compact crumpled chain organization has dimension *d* = 3 [31, 47]. In our experiments, we observe a scaling exponent of 1/*d* = 0.31 ± 0.07 for genomic separations up to *s* = 190 kb, consistent with the crumpled chain model (Fig. 2b). The smaller-than-expected average distances observed for the largest separations (*s* = 595 kb, 3.3 Mb) are most likely affected by the average folding of the chromosome [48].

**Figure 2.**
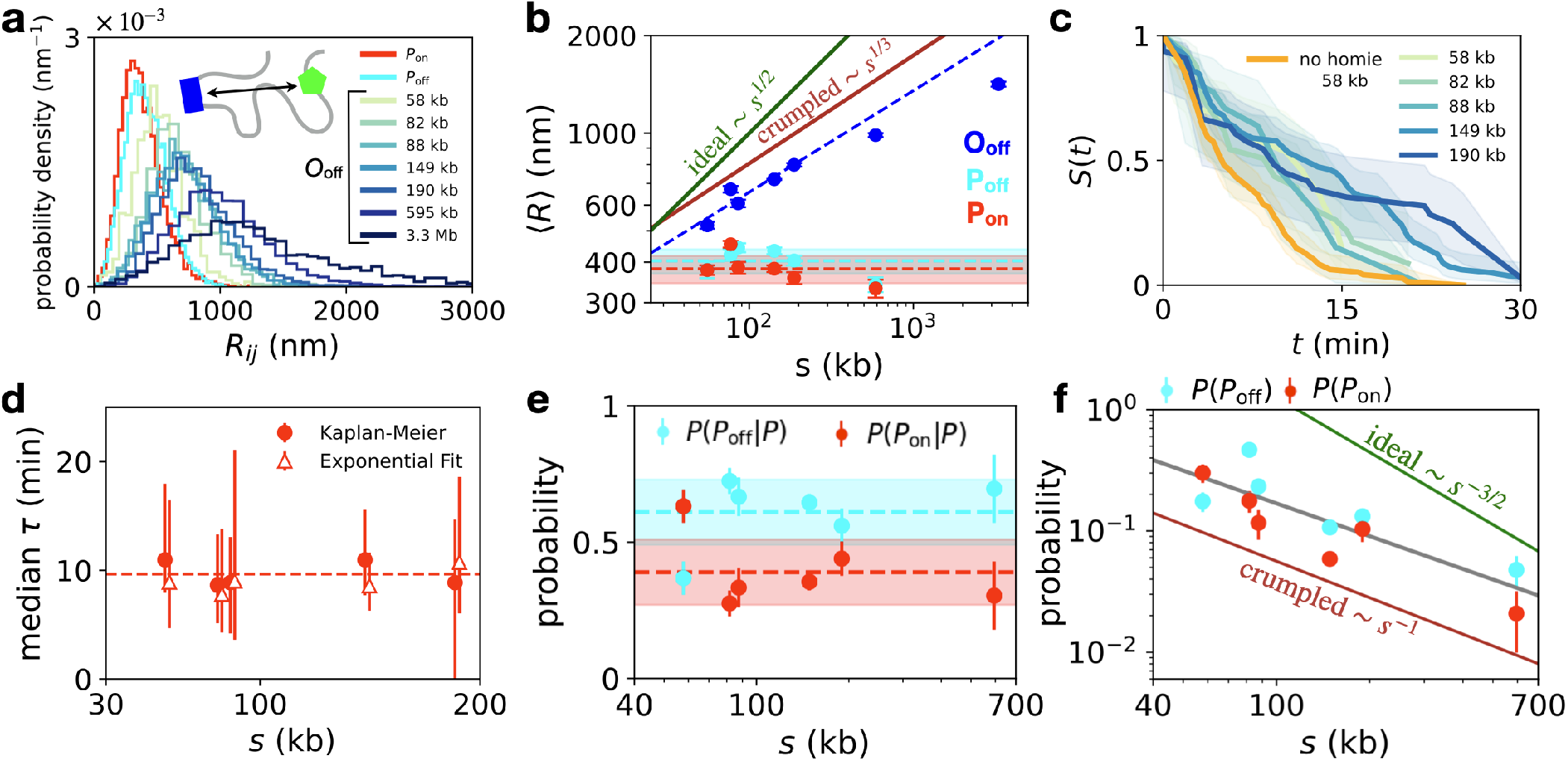
Scaling of interlocus distances and transcriptional activity across genomic separations. **a**, Probability distributions of the inter-locus distances R. Distributions are separated by state, with paired states pooled across genomic separations, and individual distributions are shown for the open state. **b**, Average inter-locus distances 〈*R*〉 for each of the three transcriptional states. Blue dashed line indicates a linear best fit to the O_off_ data for the range of genomic separations 58-190 kb, with exponent 1/*d* = 0.31 ± 0.07. Dashed cyan and red lines are average values of the interlocus distances of the P_off_ and Pon states, respectively, with shaded areas indicating error of the mean. Solid dark green and red lines indicate predictions for ideal and crumpled polymers, respectively. **c**, Survival curves *S*(*t*) of the transcriptionally active state P_on_, giving the probability that transcription remains active after time *t*. Orange curve: data for no-homie constructs (*s* = 58 kb). Curves are estimated using the Kaplan-Meier estimator which accounts for censoring, which occurs if the trajectory begins or ends in the transcriptionally active state [44]; shaded areas show 95% confidence intervals (Supplementary Section S2.4). **d**, Median lifetime of the transcriptionally active state Pon as a function of genomic separation, using the Kaplan-Meier estimator (dots) and a maximum-likelihood estimator assuming exponential decay of the survival curves (triangles; Supplementary Section S2.4). **e**, Probability of the paired on and off states conditioned on the system being in one of these two paired configurations. **f**, Overall probability of the paired configurations P_off_ and P_on_ as a function of genomic separation. Grey line: best fit with exponent 0.9 ± 0.2. Green and dark red lines indicate predicted exponents for the contact probabilities of the ideal and crumpled chain polymer models.

The distances of the paired configurations are independent of genomic separation, as anticipated, and exhibit typical distances of 350 – 400nm (Fig. 2b), consistent with previous measurements of distances within the *eve* - locus [8, 49]. Together, these results reveal a compact crumpled chain architecture of chromosome configurations in a range of genomic separations consistent with Hi-C experiments in *Drosophila* [17].

## Transcriptional activity scales with genomic separation

From the latent state trajectories revealed by our inference approach, we estimate the survival curves of the transcriptionally active state (Fig. 2c). We find a median transcriptional lifetime independent of genomic separation of (10 ± 5) min (error: std. across separations; Fig. 2d). This corresponds to about 3–5 independent rounds of transcription on average, given the typical promoter switching correlation time of the system [50]. Similarly, the relative proportion of transcriptionally active states within the paired subpopulation is insensitive to genomic separation (Fig. 2e).

In contrast, the overall probability of observing either of the paired configurations strongly decreases with genomic separation, and exhibits a power-law scaling *P*(*s*) ~ *s*^−*f*^, with *f* = 0.9±0.2 (Fig. 2f). Since transcriptional lifetimes are independent of distance, the scaling of *P*(*s*) is likely dominated by the search of the two loci to come into contact. Importantly, different polymer models make distinct predictions of the scaling of contact probabilities [22, 31, 51]: for ideal chains, *f* = 3/2, while crumpled chains exhibit *f* ≈ 1.15 [52], which is close to the scaling we observe.

To test how these results depend on the nature of the *homie* insulator-mediated focal contacts in our system, we employed a reporter construct in which the *homie* sequence is replaced by a λ DNA sequence of the same length. Interestingly, at 58 kb separation transcriptional encounters still occur, albeit with a shorter median lifetime of (4.9 ± 1.2) min. Furthermore, the probability of observing a transcriptional state is reduced from (30 ± 5)% for the *homie* version to (8.5 ± 0.8)% in the no-*homie* version. In contrast, barely any of such encounters are found for a 149 kb no-homie separation [8], where contact probability decreases from (6±1)% to > 1% when the *homie* sequence is replaced by λ DNA.

Together, these results demonstrate quantitatively how both genomic sequence and genomic separation control the rate of transcriptional encounters. Notably, the scaling of transcription probabilities with separation suggests that the transition from the open to the paired configuration is a key limiting step in transcriptional activation of distal DNA loci, which is limited by the time taken to diffuse into proximity.

## Characterizing the subdiffusive locus search process

To understand these diffusive time scales, we consider the real-time dynamics of the blue and green-labeled DNA loci. Interestingly, we find that the majority of single-cell trajectories sample the whole range of physical distances in each topological state, as they show a similar spread as the ensemble-averaged distribution (Fig. 3a-c, S8). Thus, rather than existing in constrained configurations as observed in other genomic contexts [40], this observation supports the picture of a dynamic search process exploring a broad range of distances.

**Figure 3.**
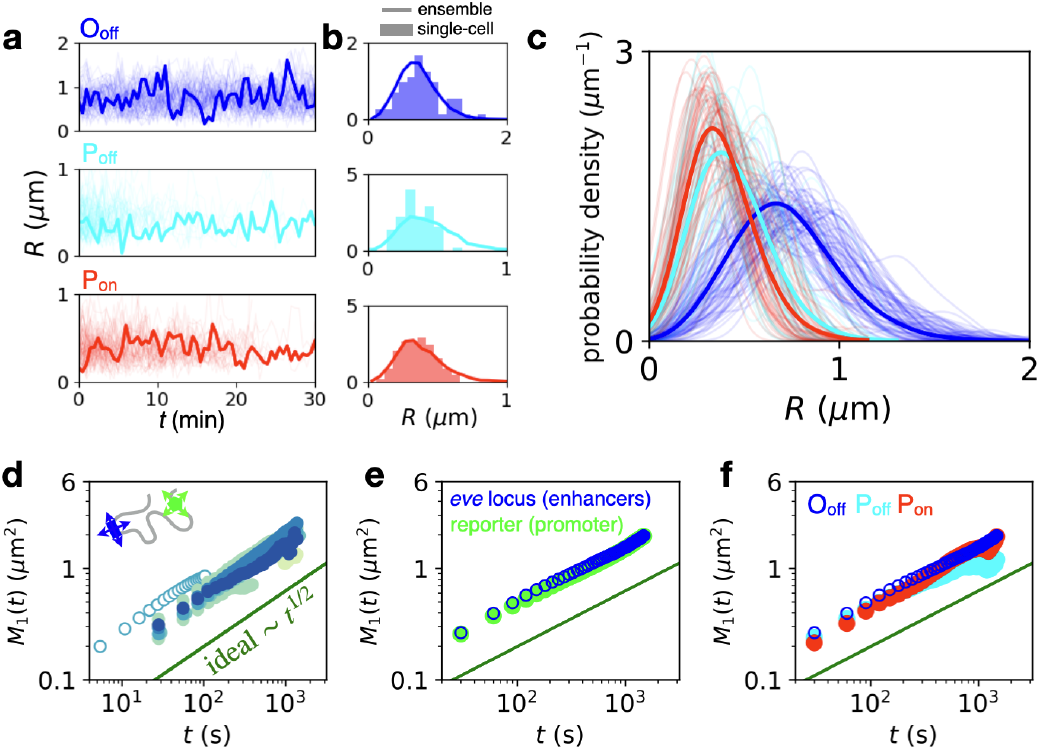
Dynamics of DNA locus search and single-locus fluctuations. **a**, Single-cell inter-locus distance trajectories for the three topological states (*s* = 149 kb). For each state, 80 trajectories are shown, with one sample trajectory highlighted in bold. **b**, Distance distributions of the highlighted trajectory in panel c (bar histogram) compared to the ensemble distribution obtained by averaging over all cells (line). **c**, Single-cell inter-locus distance distributions of all trajectories in panel c (thin lines) for the three states compared to ensemble distributions in bold (bold lines) (*s* = 149 kb). Distributions are smoothed using Gaussian kernel density estimation with a width of 100 nm. Only trajectories with at least ten time points are included to ensure sufficient statistics for comparison. **d**, Single-locus MSDs for all genomic separations (color code corresponds to Fig. 2a). Single-locus MSDs are calculated by estimating 3D MSDs from motion-corrected trajectories in the *xy*-plane of the system (Supplementary Section S3). Open data points correspond to a shorter imaging time interval Δ*t* = 5.4s (*s* = 149 kb). **e**, Single-locus MSDs comparing enhancer (blue) and promoter (green) fluctuations (*s* = 149 kb). **f**, Single-locus MSDs comparing fluctuations in the three states (*s* = 149 kb).

**Figure 4.**
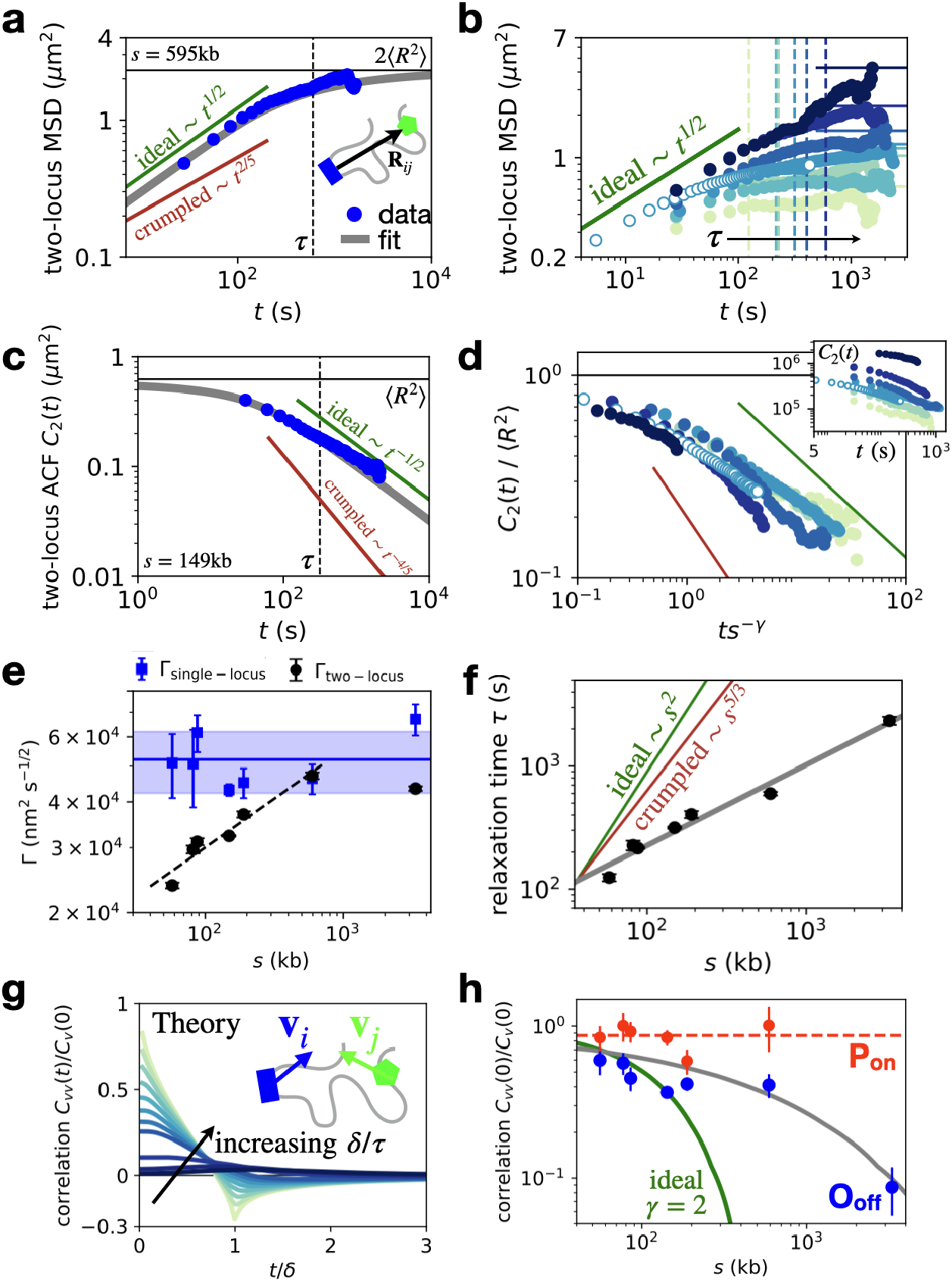
Joint dynamics of DNA locus pairs. **a**, Ideal chain Rouse prediction of the two-locus MSD 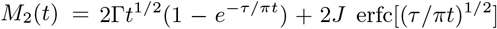 [26] (grey line), using best fit values Γ, *J*, *β* = 1/2, and *τ* = (*J*/Γ)^2^; compared to experiment (*s* = 595 kb). Green and red lines give expected scaling *t^β^* for *t* ≪ *τ* for the generalized Rouse model for ideal and crumpled chains (Supplementary Section S5). **b**, All experimental two-locus MSDs with relaxation times (dashed lines) and expected asymptotes 2〈*R*^2^〉 (solid lines; color code corresponds to Fig. 2a). **c**, Scaling of the diffusion coefficients Γ from two-locus MSD fits (black dots), compared to single-locus diffusion coefficients obtained from single-locus MSDs (Fig. 3f-h). Dashed line: best fit to two-locus diffusivity with exponent 0.27±0.03 (*s* = 58 – 595 kb); solid lines: average value of single locus diffusivities, shaded area shows error (std. calculated from total variance across separations). **d**, Two-locus autocorrelation function (ACF) *C*_2_(*t*) = 〈**R**(*t*_0_) · **R**(*t*_0_ + *t*)〉_*t*_0__ = 〈*R*^2^〉 ‒ *M*_2_(*t*)/2 (grey) compared to data (*s*=149 kb). Green and red curves indicate the power-law exponent λ = 2(1 – *d*)/(2 + *d*) of the correlation function *C*_2_(*t*) ~ *t*^λ^ for ideal and crumpled chains for *t* ≫ *τ*, respectively [39]. **e**, Collapsed correlations 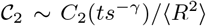 with *γ* = 0.7. *Inset*: raw correlations *C*_2_(*t*) for varying genomic separation. Open data points correspond to data obtained with a higher sampling rate. **f**, Scaling of inferred relaxation times compared to predicted ideal and crumpled chain exponents. Grey line: best fit with exponent *γ* = 0.7 ± 0.2. **g**, Predicted velocity cross-correlation functions 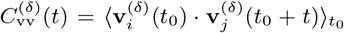 for increasing values of the dimensionless ratio *δ*/*τ* [54]. Velocities are calculated on a time-interval *δ* as **v**^(*δ*)^(*t*) = (**x**(*t* + *δ*) – **x**(*t*))/*δ*. **h**, Scaling of the zero-time velocity cross-correlation intercept normalized by the zero-time auto-correlation, 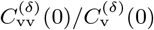, for the O_off_ (blue) and P_on_ (red) states; δ = 300s. Green line: prediction based on ideal chain Rouse scaling of the relaxation times (*γ* = 2) with an intercept determined based on the 58 kb data-point; grey line: parameter-free prediction using the inferred anomalous relaxation time scaling (*γ* ≈ 0.7) (Supplementary Section S4.3); dashed red line: average correlation in the P_on_ state.

We quantify how this search process is reflected in the motion of individual DNA loci by computing the singlelocus 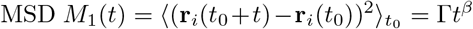, where **r**_*i*_(*t*) is the 3D position of the locus, Γ the diffusivity, and *β* the dynamic exponent. This exponent quantifies how locus diffusion scales with time and can be related theoretically to the packing of the chromosome via the fractal dimension *d*: *β* = 2/(2 + *d*) [37, 39, 53]. While the ideal chain model predicts *β* = 1/2 [33], we expect *β* = 2/5 for a crumpled polymer [37]. Our system shows a scaling exponent of *β* = 0.52 ± 0.04 across genomic separations (error bar: std. calculated from total variance across separations), for both the endogenous *eve* locus (blue) and for the ectopic reporter (green), which is close to the prediction of the ideal chain model, and consistent with previous works [26, 27, 35] (Fig. 3d,e). Notably, our data further indicate that the single-locus dynamics are not affected by transcriptional activity, unlike previous accounts [42], as they are consistent across the three topological states (Fig. 3f).

To further understand how the locus dynamics are determined by the interplay of chromosome organization and single-locus dynamics, we analyze the joint dynamics of the two coupled chromosomal loci. From the statistics of the 3D distance vector **R**(*t*), we compute the two-locus MSD *M*_2_(*t*) = 〈(**R**(*t*_0_ + *t*) – **R**(*t*_0_))^2^〉_*t*_0__ [26], which quantifies the crossover between two intuitive regimes. While at small time-lags, the MSD is determined by the independent diffusion of the two loci (*M*_2_(*t*) = 2Γ*t*^*β*^), it exhibits a cross-over to a plateau at large times, given by the average squared inter-locus distance (*M*_2_(*t*) = 2〈*R*^2^〉) (Fig. **??**a,b). Consistent with the observed single-locus dynamics, we find that also the small-time limit of the experimental two-locus MSDs exhibits an exponent close to the ideal chain with 1/2 (Fig. **??**a). Similarly, for large time-lags, the two-locus auto-correlation reveals agreement with the ideal chain scaling (Fig. **??**c,d). Thus, the full time-dependence of the MSD is well described by the ideal chain predictions, both for single and coupled loci.

## Inter-locus relaxation times exhibit an anomalous scaling with genomic separation

Having established the static and dynamic properties of the system, we now ask about the consequences of these features for the time scales of the two-locus search process. This process is determined by the interplay of chromosome dynamics and organization and can be characterized by a relaxation time *τ*, which corresponds to the time scale of the crossover of the two regimes of the two-locus MSD (Fig. **??**a). Specifically, τ is the time taken by the two loci to diffuse (dynamics) over their typical distance of separation (organization): Γ*τ*^*β*^ ~ *s*^2/*d*^. This relationship predicts a scaling of relaxation times with genomic separation *τ* ~ *s^*γ*^*: for ideal chains with fractal dimension *d* = 2 and a diffusion exponent *β* = 1/2, this yields the classical result *γ* = 2. In contrast, for crumpled chains, *β* = 2/5 and *d* =3, yielding *γ* = 5/3.

To infer the relaxation time in our data as a function of genomic separation, we perform a Bayesian fitting of the two-locus MSD with the ideal chain expression [26] (Supplementary Section S4). Interestingly, we find that the fitted two-locus diffusion coefficient increases with genomic separation up to 595 kb, with an approximate scaling Γ(*s*) ~ *s*^0.27±0.03^ (Fig. **??**e). This scaling appears to plateau for the largest genomic separation (3.3Mb) at a value close to the single locus diffusion, which remains approximately constant across separations (Fig. **??**e). Notably, the absolute value of the diffusivity at the plateau is almost 20-fold larger than previous measurements in mammalian stem cells with similar genomic separation [26], suggesting comparatively fast chromosome dynamics (Fig. S23).

To infer the relaxation time as a function of genomic separation, we combine our estimate of the two-locus diffusivity with the average inter-locus distances. Due to the combination of static and dynamic exponents in our system, as well as the scale-dependent diffusivity, we find an anomalous relaxation time scaling with an exponent *γ* = 0.7 ± 0.2 (Fig. **??**f). This exponent corresponds to a much shallower scaling with separation than predicted by either the ideal or crumpled chain theory. This result is further confirmed by a data collapse of the two-locus auto-correlation functions (Fig. **??**d, Fig. S20). While these results are derived from the trajectories in the O_off_ state, they are insensitive to the details of the state inference (Fig. S11). In sum, the key result here is that the relaxation time, which sets the time scale of two-locus encounters is much less dependent on genomic separation than predicted by existing polymer models.

## Anomalous relaxation time scaling induces long-ranged velocity correlations

The anomalous relaxation time scaling makes a key prediction for the correlations of the absolute motion of DNA loci, quantified by the velocity cross-correlation 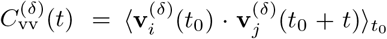. These correlations are determined by the relaxation time through the dimensionless ratio *δ*/*τ*, where *δ* is the experimental observation time-scale (Fig. **??**g) [54]. Having determined the relaxation times *τ*, we can therefore make a parameter-free prediction of the correlations, which decay significantly more slowly than for the ideal Rouse model (Fig. **??**h, green and grey lines). Notably, we find that the experimental correlations are quantitatively captured by this parameter-free prediction (Fig. **??**h), including the full time-dependence of the correlations (Fig. S22). This demonstrates that the anomalous relaxation time scaling indeed leads to long-ranged velocity cross-correlations of chromosomal loci, pointing towards potential long-range interactions.

## Discussion

We developed an experimental approach to perform *in vivo* imaging of the joint dynamics of enhancer–promoter pairs with varying genomic separation and simultaneous monitoring of their transcriptional output. Observing the dynamics of pairs of DNA loci has only become possible recently and has been done for tagged DNA loci at a single fixed genomic separation [8, 26, 27, 36]. Here, we show how imaging across genomic separations gives insight into the relative motion, dynamic encounters, and transcriptional activation of such loci.

Many features of the two-locus dynamics, including the subdiffusive exponent close to 0.5, are very well conserved with measurements of CTCF sites at TAD boundaries in mammalian systems [26, 27], despite CTCF not being essential for Drosophila embryogenesis [55]. In absolute numbers, however, our measurements reveal strikingly large diffusion coefficients of DNA loci, around 20-fold larger than in mammalian cells [26] (Fig. S23). Early fly development follows an extremely tight schedule, suggesting that the chromosome dynamics may have evolved to operate on much faster time scales than mammalian systems. In contrast, the median lifetime of focal contacts in our system of (12 ± 5) min is well within the range of typical CTCF loop lifetimes of 10 – 30 min in mammalian cells [26, 27]. These time scales facilitate transcriptional lifetimes of (10 ± 5) min in our system, which in the absence of the homie insulator are reduced to (4.9 ± 1.2) min, highlighting the importance of focal elements for contact formation in *Drosophila*.

To initiate such transcriptional encounters, the two loci must diffuse into physical proximity, at a time scale set by the relaxation time. Indeed, we find that the lifetimes of the unpaired O_off_ state correlate well with the relaxation times, but are approximately 10 times larger on average (Fig. S16). While the absolute values of these unpaired lifetimes depend on the biochemical properties of the focal elements, the relaxation time sets a lower bound and determines the dependence on genomic separation.

We demonstrate how key features of our system – tight crumpled chain packing, subdiffusion with exponent 0.5, and a separation-dependent two-locus diffusivity – lead to relaxation times that are much less dependent on genomic separation than predicted by existing polymer models. Indeed, for an ideal Rouse polymer, the relaxation time for our largest genomic separation (3.3 Mb) would be ~ 3000 times longer than for the shortest 58 kb separation. Our measurements however reveal that it only takes ~ 20 times longer, corresponding to a more than 100-fold reduction. This reduced dependence on distance implies that the probability of reaching and maintaining the spatial proximity required for transcription is similar for enhancers dispersed across the chromosome, allowing them to find their target promoter efficiently. This might be one of the reasons for why evolution can act on distal sequences from a given target promoter. Overall, our findings have crucial implications for the spatiotemporal organization of the cell nucleus, including the dynamics of long-range focal contacts [28] and mammalian enhancer-promoter interactions [9–12, 43].

From a polymer physics perspective, our measured exponents suggest that the relationship between static and dynamic properties in the generalized Rouse framework, which relies on the assumption of local friction, does not apply to chromosomes. This suggests that long-range interactions, such as hydrodynamics or active motor-mediated interactions [56, 57] could play a role. Indeed, the simplest polymer model that relaxes the Rouse assumption and includes long-range hydrodynamic interactions, the Zimm model [33], predicts a scaling relationship of relaxation times with genomic separations with an exponent of *γ* = 1, which is close to our measured value of *γ* ≈ 0.7. Furthermore, the observed separation-dependent diffusivity points to additional heterogeneities along the polymer. Such heterogeneities could be due to a number of processes, such as cross-linking [40], out-of-equilibrium activity [57], entanglements [58], or the presence of condensates [18–20]. Together, these processes may orchestrate the anomalous scaling of relaxation times with genomic separation. In future work, the mechanistic underpinnings of our findings should be tested through polymer simulations [32, 40, 51, 59–64] to generate hypotheses for new sets of experiments.

## Acknowledgements

We thank S. Grosse-Holz, L. Mirny, G. Tkacik, R. Evearers, L. Giorgetti, A. Grosberg, A. Spakowitz, M. Levo, A. Rosa, and V. Scolari for helpful discussions. We thank K. Bystricky for introducing us to the ParB/parS system, and F. Payre and P. Valenti for sharing a ParB-eGFP plasmid and the parS sequence. This work was supported in part by the U.S. National Science Foundation, through the Center for the Physics of Biological Function (PHY-1734030), and by National Institutes of Health Grants R01GM097275, U01DA047730, and U01DK127429. D.B.B. was supported by the NOMIS foundation as a NOMIS Fellow and by an EMBO Postdoctoral Fellowship (ALTF 343-2022). H.C. was supported by the Charles H. Revson Biomedical Science Fellowship.

