## Supplementary Material for "Stochastic motion and transcriptional dynamics of pairs of distal DNA loci on a compacted chromosome"

### Materials and Methods

**1 Experimental methods** **2**

**5 Two-locus dynamics: theory** **32**

**6 Comparison to mESC data** **35**

**7 Data and materials availability** **36**

**8 Movie description** **36**

### 1 Experimental methods

#### 1.1 Generation of the distance series

The endogenous *eve* locus was tagged with MS2 stem loops. First, an *attP* site was integrated at the first intron of *eve* using CRISPR-mediated homology-directed repair [1]. The two Cas9 cutting sites are marked in Fig. S1a. Second, an attB-MS2 plasmid was used to deliver MS2 into the attP site. A genomic source of phiC31 integrase (BDSC #34770) was used for the second injection. The synthetic eve-MS2 allele carries 10.6 kb between the two Cas9 cutting sites.

A series of attB landing site lines were established by mobilizing the P-element construct P-2xattB-mW from a CyO balancer to the chromosome containing the eve-MS2 allele. A Δ2-3 P-transposes line (BDSC #3629) was used for the transposition. Offspring displaying both Cy+ and w+ were subject to inverse PCR to map the location of the attB insertions. attB insertions at -3321.9, -589.3, +47.5, +77.5 and +179.3 kb from the eve promoter in cis are used in this study. Furthermore, in order to avoid the insertion biases of P-element, two MiMIC lines (MI02916 and MI00239, [2]) were recombined with the eve-MS2 chromosome to yield the -75.9 and -186.8 kb lines. The exact insertion points of these lines and their conversion to distances between the blue and green labelled spots are listed in Table S1.

Finally, the PP7 reporter transgenes (parS-homie-evePr-PP7 or parS-lambda-evePr-PP7) were integrated into the landing sites described above through recombination-mediated cassette exchange using BDSC #34770 as the integrase source. Transgenic flies with insertion orientations shown in Fig. S1a were obtained and used for the imaging experiments.

Fluorescent protein tagged MCP, PCP and ParB lines were made as previously described [3]. For maternal MCP::3xmTagBFP2 and PCP::3xmKate2, a genomic landing site at 38F1 was used. For maternal ParB::eGFP, a landing site at 89B8 was used. All microinjections were performed as described previously [4].

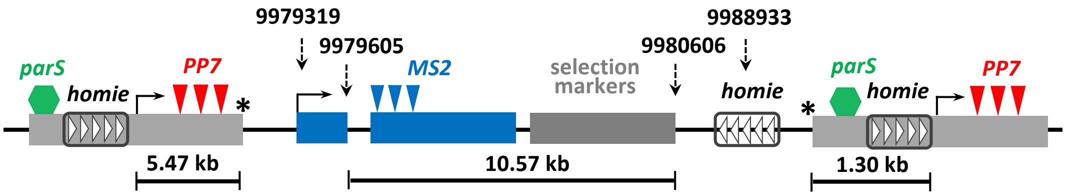

Figure S1: **Map of the genomic constructs.** The transcription start site (TSS) of *eve* (2R:9979319), the center of the endogenous *homie* element (2R:9988933) and the two Cas9 cutting sites (2R:9979605 and 2R:9980606, marked by stars) used for *MS2* knock-in are labelled with dashed arrows. Each *evePr-PP7* reporter line is named after the distance between its insertion site and the eve TSS, with a sign indicating upstream (-) or downstream (+) to the eve locus. Note that the new eve allele is larger than the endogenous one due to the presence of transgenic selection markers (GFP and RFP). The *MS2-parS* distance can be calculated from the distance shown below the map. Genomic distances and size of the sequence elements are drawn in scale.

#### 1.2 Microscopy

Virgins carrying three fluorescent protein fusions (yw; MCP::3xmTagBFP2/PCP::3xmKate2; ParB::eGFP/+) were crossed with males containing the eve-MS2 allele and the parS-evePr-PP7 reporter transgenes. Embryos from the crosses above were dechorionated manually and mounted as described in [5].

| Name | Breaking point on 2R | Distance to *eve* TSS (bp) | *MS2-parS* distance (kb) |
|---|---|---|---|
| -3322 | 6657453 | -3321866 | 3327.7 |
| -589 | 9389997 | -589322 | 595.1 |
| -142 | 9836454 | -142865 | 148.7 |
| -76 | 9903417 | -75902 | 81.7 |
| +47 | 10026855 | 47536 | 57.8 |
| +77 | 10056812 | 77493 | 87.8 |
| +179 | 10158620 | 179301 | 189.6 |

Table S1: **Details of the distance series.** All breaking point coordinates are from the Dm5 reference. Throughout the text, we cite the *MS2-parS* distances, and refer to their rounded values in legends for simplicity.

A Leica SP5 confocal microscope with an oil immersion 63x NA 1.44 objective was used in all imaging experiments. Three laser lines at 405 nm, 488 nm and 591 nm were used to excite mTagBFP2, eGFP and mKate2, respectively. For each z slice, the emission channels for 405 nm and 591 nm were acquired simultaneously, followed by the 488 nm channel. Power measured at the objective for the 405 nm, 488 nm and 591 nm lines was 0.4 $\mu$W, 1.1 $\mu$W and 0.5 $\mu$W, respectively. Three HyD detectors in photon counting mode were used to collect fluorescence emission spectra.

The voxel size for all images is 107x107x334 nm$^3$ (x/y/z). The stack interval for all time-lapse videos was 28 s, except for the ones shown in Fig. 3c, 4b, and 4d, where the stack interval is 5.4 s. For the data with a 28 s frame rate, an 8 $\mu$m z-stack covering the whole nucleus along the apical-basal direction is acquired during the 28 s time interval. Images were taken at 1,024 x 256 x 25 voxels and focused on the posterior half of the embryo, encompassing *eve* stripes 3–7. Embryos were timed by their exit from mitosis 13 [6]. All embryos were imaged from 20 $\pm$ 2 min to 55 $\pm$ 2 min in cell cycle 14. Based on a localization control construct, the measurement errors in spot localization in this setup are 180 $\pm$ 6 nm (mean $\pm$ s.e.m.) [3]).

##### 1.3 Nucleus and DNA locus segmentation and tracking

Segmentation and tracking of cell nuclei and DNA loci was done as described before [3]. Briefly, we computed the difference between the blue and red channels (since NLS::MCP::3x mTagBFP2 is enriched in the nuclear compartment while ParB-eGFP is enriched in the cytoplasm) by subtracting the maximum $z$-projection of the green channel from the blue channel, followed by Gaussian blurring ($\sigma$ = 5) and binarization (local Otsu's threshold at 5 x 5 $\mu$m$^2$ [7]). To account for the motion of the whole embryo and the nuclei during imaging, we inferred the nuclear movement by minimizing the cross-correlation between nuclear masks of two consecutive frames. All nuclear segmentation and tracking results were scrutinized manually.

For DNA locus segmentation, we first identify candidate loci using sharpened raw image stacks from each of the three channels using a 3D bandpass filter (size 11 x 11 x 7 pixels), derived by subtracting a uniform filter from a Gaussian kernel ($\sigma$ = (1, 1, 0.6) pixel). All local maxima of this filtered image are considered as candidate loci. At each local maximum, we constructed cylinder mask with diameter of 13 pixels (1.4 $\mu$m) and a height of 7 pixels (2.3 $\mu$m). We then set an intensity threshold such that the maximum number of candidate loci in the nucleus at each time point was less than 20, followed by filtering the set of candidate locus using information on nuclear lineage, spot tracking and the relative location of spot pairs.

For DNA locus tracking, we calculated the sub-pixel centroid of each candidate locus, only considering candidate loci located in the corresponding nuclear region obtained from the nuclear segmentation results. Spot tracking was then performed in three steps: a pre-tracking step, a gap-filling step and a Bayesian filtering step [3].

##### 1.4 Chromatic aberration correction

Imaging the positions of the *eve*-locus and the ectopic reporter with different wavelengths introduces chromatic aberrations, i.e. an instrumental error. This appears as a systematic error in our data (constant position displacements), which can vary across the field of view. Since positions of imaged loci are random, we assume that on average they have to be centered at zero, and any systematic errors are caused by the chromatic aberrations between the two channels. We perform a correction of this systematic error using data-driven and internally controlled (i.e., for the entire data set on any given day of data collection) calibration [3]. We consider the entire field of view and make a correction at each position in the 3D field of view.

We assume that the mean of the blue-green vector along each dimension is zero. For example, the probability that a blue spot appears above the associated green spot should be the same as the probability that it is below the associated spot. By applying linear regression to each dimension of displacements (x, y, and z) and extracting a 3x3 matrix, effectively the slope of this bias across the field of view in each dimension, the spot coordinates from a given color channel are scaled by the matrix to obtain the correction vector. This correction vector is the difference in the scaling of two channels, using one as a reference for the other to be matched to. Subtracting the correction vector from the original spot gives the corrected spot centroid, which can then be used for displacement calculation. This procedure effectively rescales the coordinates of all spots as to readjust the aberration slope across the fields of view to zero.

The assumptions underlying this approach were tested explicitly using control experiments with colocalized fluorescent proteins and three-color TetraSpec beads [3] (Fig. S2). This approach also allowed estimation of the residual measurement error in spot localization as being 180 $\pm$ 6 nm, in line with the errors inferred from our MSD analysis, see below (section 4.1).

Chromatic aberration correction was performed on a weekly basis, using all the embryos of the same genotype imaged over the week. An example of the correction is given in Fig. S2. We used this data-driven and internally controlled calibration scheme to estimate the blue-green distance for all genomic constructs, which we report in the main text.

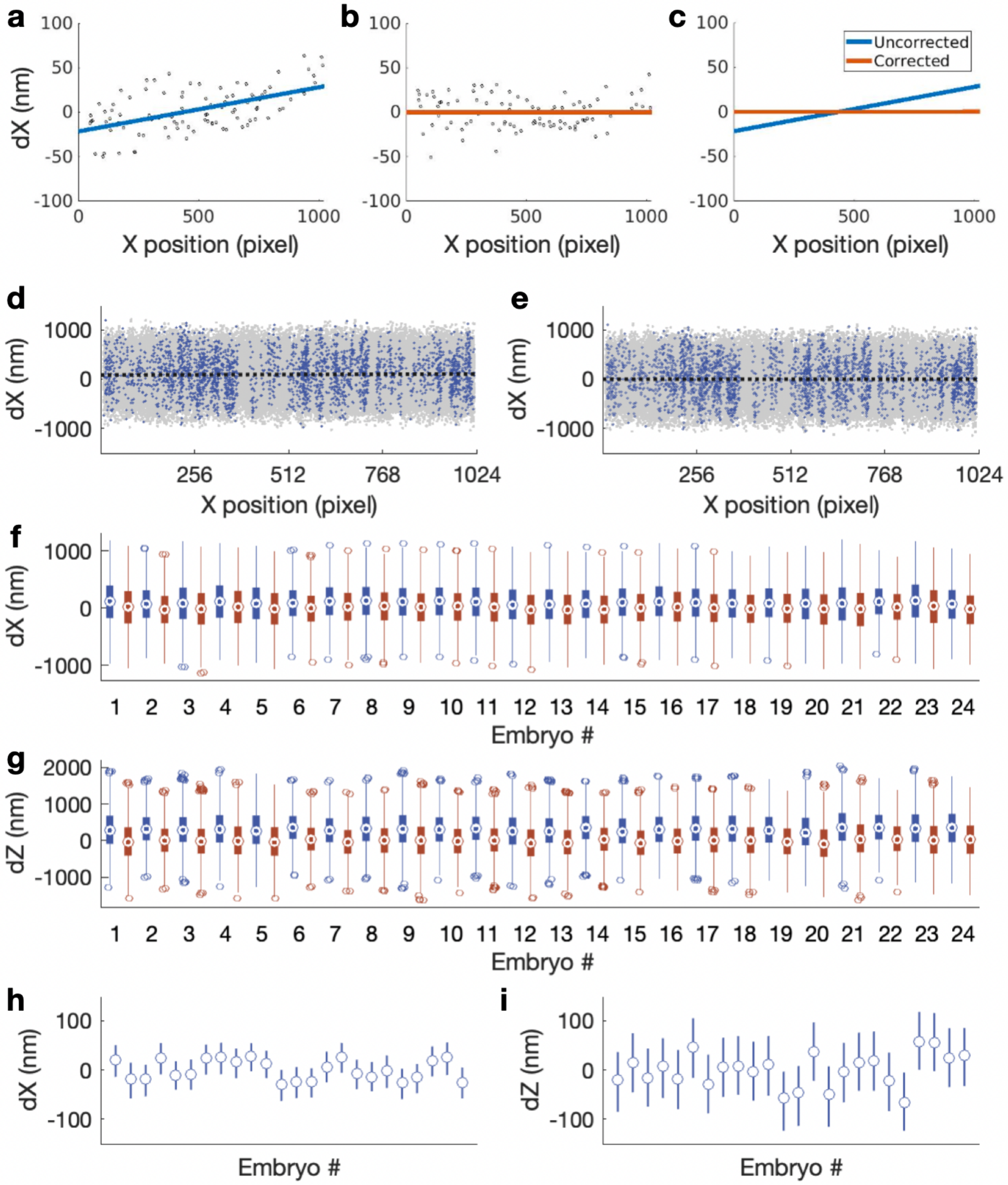

Figure S2: **Correcting chromatic aberrations.** **a-c**, Chromatic aberration distance bias before (a) and after (b) correction with comparison (c) in the *x*-component of blue and green channels of calibration beads (Tetraspeck) for one image (n = 95). **d**, A representative dataset showing raw *MS2-parS* distance, i.e. the raw distance between a blue-green spot pair (E-P distance) as a function of the spot pair position in the field of view (1 pixel = 107 nm). 24 -142 kb *homie-evePr-PP7* embryos imaged over a week are included (gray points). These embryos form a chromatic aberration correction batch. Blue data points are from a single embryo. Dotted line shows the result of the linear fitting. The slope ($0.0186 \pm 0.0045$) is significantly different from zero ($p = 3.96 \times 10^{-5}$, two-tailed t-test), indicating chromatic aberrations due to the optical setups. **e**, E-P X distance after chromatic aberration correction described in Supplementary Methods. Both the slope and intercept are zero for the fitted line after the correction. **f,g**, Boxplots showing distributions of the raw (blue boxes) and corrected (red boxes) E-P X (f) and Z (g) distance for each embryo in the correction batch. Medians are shown. Boxes represent the interquartile range, and the whiskers extend to 2.5 times the interquartile range above or below the medians. **h,i**, Estimating errors associated with the correction. The errors are estimated by obtaining a correction matrix from every single embryo and applying it to all embryos in the same batch to get a distribution of the corrected E-P distance. Means of E-P distances calculated from the correction matrix derived from different single embryos are plotted, with bars showing 2.5-97.5 percentile range for dX (h) and dZ (i). The error (standard deviation) is 21 and 35 nm for dX and dZ, respectively.

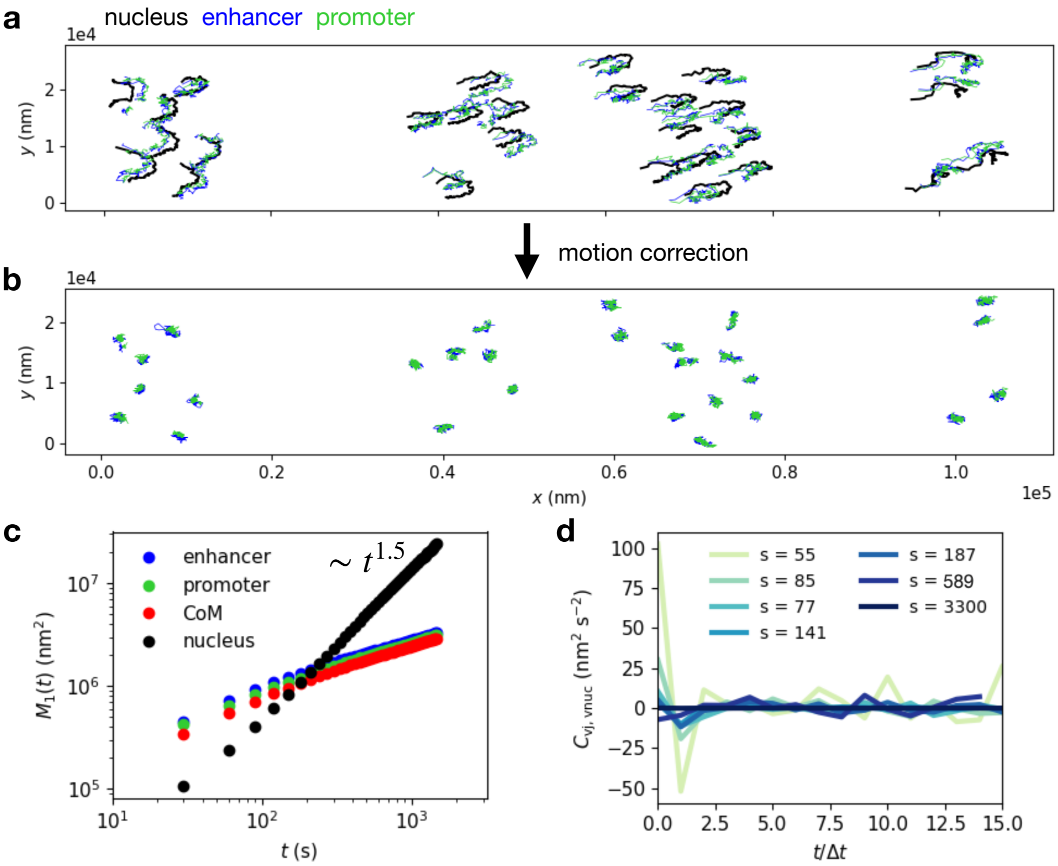

Figure S3: **Tracking of cell nuclei.** **a**, Trajectories of nuclei (black), enhancer (blue) and promoter (green) within a single embryo ($s$=149 kb) in the lab frame of reference. **b**, Motion corrected DNA locus trajectories. **c**, Single-locus MSD $M_1(t)$ (Eq. (S25)) of the enhancer (blue), promoter (green), their center of mass $(\mathbf{r}_1(t) + \mathbf{r}_2(t))/2$ (red), and the nucleus trajectories (black). Approximate scaling of the nucleus MSD scaling is indicated. **d**, Velocity cross-correlation of the motion corrected locus trajectory and the nucleus trajectory, indicating no correlation beyond the first time point $t = \Delta t$, as expected for purely random measurement errors in the nucleus tracking (section 4.3).

#### 1.5 Nucleus motion correction

Following tracking of the DNA loci, two further operations are performed: (i) we corrected for chromatic aberration to gain access to the dynamics of the 3D distance vector $\mathbf{R}_{ij}$; (ii) we performed motion correction of the trajectories to obtain the absolute motion of DNA loci within the nucleus frame of reference.

To correct chromatic aberration, we perform a data-driven, internally controlled calibration; and approach that was tested extensively in previous work [3]. Briefly, we pooled raw instantaneous locus-pair distances across time, nuclei, and embryos, and analyzed the raw distances as a function of the locus-pair positions in the image field of view. To correct aberration, we applied a multivariate normal regression model to infer the correction matrix that allows the calculation of calibrated distances (see section above), which were used throughout further analysis.

To perform motion correction, we subtract the nucleus trajectories from the DNA locus trajectories (Fig. S3a,b). In many experimental systems, this is challenging as nuclei often undergo complex combinations of rotation, shape changes, and translation. During the final phase of the syncytial blastoderm stage of the developing *Drosophila* embryo, nuclei undergo very little relative motion, do not rotate, and simply show affine global translation, which we can track as described above. As the same nucleus trajectory is subtracted from both DNA loci, measurement errors in the nucleus tracking could introduce spurious correlations in the observed relative motion of the two DNA loci. To exclude systematic measurement errors in the nucleus tracking, we calculate the velocity cross-correlation of the motion corrected locus trajectory and the nucleus trajectory (Fig. S3d), which is consistent with the expectation for time-uncorrelated measurement errors in nucleus tracking (section 4.3). Furthermore, we find no traces of spurious motion in the trajectories of the DNA locus center of mass, as its MSD follows the same subdiffusive exponent as that of individual loci, which should be contrasted with the superdiffusive exponent of the nucleus movement (Fig. S3c). Together, these tests demonstrate that nucleus tracking and motion correction only introduces random time-uncorrelated measurement errors in the locus trajectories, which we account for using error-correction in the velocity cross-correlations (section 4.3).

### 2 Topological and transcriptional state analysis

#### 2.1 Hidden Markov Model inference

Based on the architecture of our genetic system comprising the enhancer and promoter, which are each flanked by a homie insulator element, the system can be in one of three states: (i) A state where the two homie elements are unbound, termed $O_{\text{off}}$. In this state, we expect large inter-locus distances. (ii) A topologically distinct state where the two homie elements are bound, but where transcription is inactive, termed $P_{\text{off}}$. In this state, we expect small inter-locus distances, and no signal in the red channel. (iii) A state where the two homie elements are bound and transcription is active, termed $P_{\text{on}}$. In this state, we expect small inter-locus distances, and the red channel to be on. The existence of these three states was demonstrated previously using various control experiments with individual elements of the genetic cassette missing [3]. To verify the assumption that active transcription corresponds to smaller physical distances in all sampled genomic separations, we plot interlocus distances depending on the intensity of the red channel (Fig. S4). This confirms that transcriptionally active states are associated with smaller physical distances, consistent with previous observations [3].

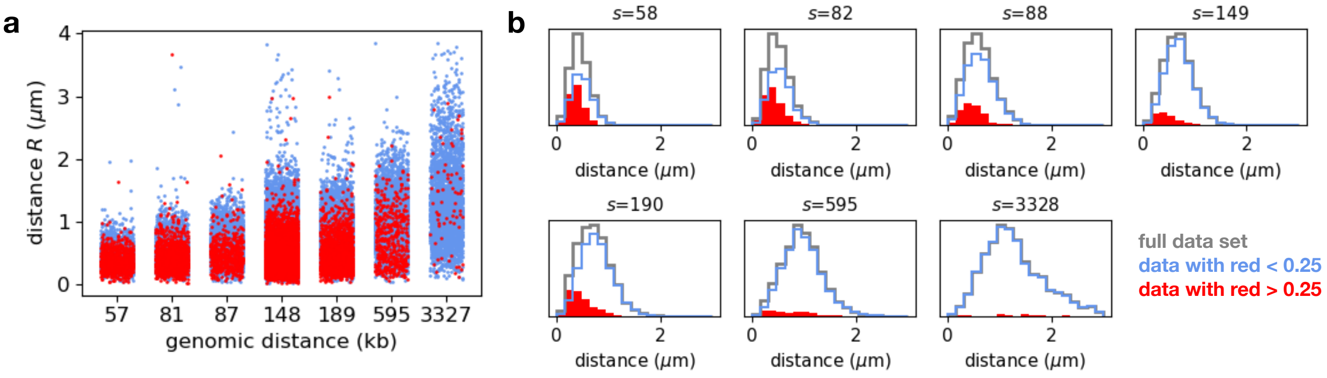

Figure S4: **Interlocus distances as a function of transcriptional state. a**, Scatter plots of the entire data set for each genomic separations, with points colored by whether the red signal is on (red intensity $> 0.25$; red), or off (red intensity $< 0.25$; blue). **b**, Histograms of the interlocus distances for the entire data set (grey), points where the red signal is on (red intensity $> 0.25$; red), or off (red intensity $< 0.25$; blue).

Together, these arguments therefore suggest an underlying structure of state transitions as shown in Fig. S5a. Transitions between $O$ and $P$ toggle between two distinct topological states; transitions between OFF and ON toggle between two transcriptionally distinct states. Importantly, the details of the state inference performed here does not influence our later analyses on the polymer configurations or dynamics, which are based directly on the observed distance trajectories, which do not sensitively depend on the inferred underlying states (see Supplementary Section 2.6).

To infer the particular state of the system at each moment in time, we modeled the data (inter-locus distance and intensity time-series) as a Hidden Markov Model (HMM) [8] calibrated using a Markov Chain Monte Carlo (MCMC) sampler [9]. The general idea underlying 'classical' inference is to maximize the probability of the data under some model, namely to find the parameters $\Theta$ that maximize the likelihood of the data $\mathcal{D}$. In this manuscript we adopted a Bayesian approach, simultaneously estimating the state identity and the probability of the state transition rates given the observed data, i.e., the joint posterior distribution $P(\Theta|\mathcal{D})$, using Bayes' rule:

$$P(\Theta|\mathcal{D}) = \frac{P(\mathcal{D}|\Theta)P(\Theta)}{P(\mathcal{D}) \equiv \int P(\mathcal{D}|\Theta)P(\Theta)d\Theta}, \tag{S1}$$

where $P(\Theta)$ is the prior that encodes for prior knowledge about the parameter values, which we take to be log-uniform here since these are real positive and have some unknown scale [10].

The likelihood of the data $P(\mathcal{D}|\Theta)$ is expressed in terms of the likelihood of the individual nuclei time-series, which are independent:

$$P(\mathcal{D}|\Theta) = \prod_{i=1}^{N_{\mathcal{D}}} P(\mathcal{D}_i|\Theta), \tag{S2}$$

where $\mathcal{D}_i = \{(R_0, R_1, ...), (I_0, I_1, ...)\}$ corresponds to the time series of the inter-locus distance $R$ and the transcriptional fluorescence intensity $I$ for a single nucleus. Assuming a HMM, the likelihood for a single nucleus takes the following form:

$$P(\mathcal{D}_i|\Theta) = \sum_{\{\Lambda\}} \left[ \prod_{t=1}^{N_t} P(R_t, I_t|\lambda_t) P(\lambda_t|\lambda_{t-1}) P(R_0, I_0|\lambda_0) P(\lambda_0) \right], \tag{S3}$$

where $\lambda_t$ is the hidden state of the system at time $t$ (either $O_{\text{off}}$, $P_{\text{off}}$, or $P_{\text{on}}$, Fig. S5a), $P(R_t, I_t|\lambda_t)$ the "emission" probability, $P(\lambda_t|\lambda_{t-1})$ the transition probability, $P(\lambda_0)$ the initial state probability, and the sum is performed over all possible state configurations $\Lambda \equiv (\lambda_0, \lambda_1, ...)$. The likelihood above is computed efficiently using the forward algorithm [8].

For our specific case, we modeled the emission probability that relates the data to the hidden states, as two independent probability distributions, i.e. $P(R, I|\lambda) = P(R|\lambda)P(I|\lambda)$. The first is the probability distribution $P(R|\lambda)$ for the inter-locus distance $R_{ij}$ given each state $\lambda$. We find that the Maxwell-Boltzmann distribution provides a good fit to the experimental distributions of inter-locus distances (Fig. S5e). We therefore assume this general functional form for the distance distributions:

$$P(R_{ij}|\sigma_R) = 4\pi R_{ij}^2 \left( \frac{3}{2\pi\sigma_R^2} \right)^{3/2} \exp\left( \frac{-3R_{ij}^2}{2\sigma_R^2} \right). \tag{S4}$$

This distribution is quantified by a single parameter $\sigma_R$, which is the average distance, i.e. $\langle R_{ij} \rangle = \sigma_R$. This is the probability distribution of end-to-end distances of an ideal chain polymer, with $\sigma_R^2 = Na^2$, where $N$ is the number of monomers and $a$ is the monomer length [11]. We choose this general functional form here as it simplifies the inference (single parameter) and also provides a good fit to our empirical observations.

The second distribution $P(I|\lambda)$ models the fluorescence $I$ of the red channel (as a measure of transcriptional activity) conditioned on whether the state $\lambda$ is transcriptionally active ($P_{\text{on}}$) or inactive ($O_{\text{off}}$ or $P_{\text{off}}$). To capture the broad distribution of fluorescence intensities in the transcriptionally active state ($P_{\text{on}}$), we employ a Rayleigh distribution as an effective model to approximate the expected overdispersed Poisson noise due to transcriptional bursts and the additive Gaussian measurement noise :

$$P(I|\sigma_I) = \frac{I}{\sigma_I^2} \exp\left(\frac{-I^2}{2\pi\sigma_I^2}\right). \tag{S5}$$

Importantly, this distribution remains simple as it is quantified by a single parameter $\sigma_I$, and turns out to be flexible enough to also account for the distribution of background fluorescence intensities in the transcriptionally inactive state ($O_{\text{off}}$ or $P_{\text{off}}$). Indeed, the above distribution captures the empirical distribution of intensities decently well (Fig. S5f), and the specific choice we made here did not impact significantly our state calling.

Next, the transition between the three states ($O_{\text{off}}$, $P_{\text{off}}$, and $P_{\text{on}}$) are defined by the generator or Laplacian matrix of the system according to the network in Fig. S5a:

$$G = \begin{pmatrix} -f_1 & b_1 & b_3 \\ f_1 & -b_1 - f_2 & b_2 \\ 0 & f_2 & -b_2 - b_3 \end{pmatrix}. \tag{S6}$$

The resulting transition probabilities $P(\lambda_t|\lambda_{t-1})$ in Eq. S3 are then computed as the matrix exponential of $G$, i.e. $P(\lambda_t|\lambda_{t-1}) \equiv \exp[G\Delta t]$, where $\Delta t$ is the sampling time interval. The initial state distribution $P(\lambda_0)$ is chosen as the steady state distribution of the system, which is obtained by solving the linear system $GP = 0$.

Lastly, $\Theta$ represents the entire set of parameters governing the observed distances, the observed intensities and the state transition rates with a total of six physical scales and five kinetic parameters that need to be inferred:

$$\Theta = \{\sigma_R(O_{\text{off}}), \sigma_R(P_{\text{off}}), \sigma_R(P_{\text{on}}), \sigma_I(O_{\text{off}}), \sigma_I(P_{\text{off}}), \sigma_I(P_{\text{on}}) f_1, f_2, b_1, b_2, b_3\}. \tag{S7}$$

Due to the underlying symmetries of our model, the eleven parameters above are not fully identifiable (structural identifiability). Indeed, in absence of structure imposed on the $\sigma_R$ and $\sigma_I$, the states $O_{\text{off}}$, $P_{\text{off}}$ and $P_{\text{on}}$ could be permuted and the inference would be degenerate. However, as we mentioned in the beginning of this section (see Fig. S4), we have some prior knowledge about the structure of the system. From the data, it is clear that $\sigma_R(O_{\text{off}}) \geq \sigma_R(P_{\text{off}}) \simeq \sigma_R(P_{\text{on}})$ and $\sigma_I(O_{\text{off}}) \simeq \sigma_I(P_{\text{off}}) \leq \sigma_I(P_{\text{on}})$ should be satisfied. Thus, to break the symmetries, we decided to enforce $\sigma_R(P_{\text{off}}) = \sigma_R(P_{\text{on}}) \equiv \sigma_R(P)$, effectively reducing the number of parameters by one. The same treatment could be performed on the $\sigma_I$'s (i.e., enforcing $\sigma_I(O_{\text{off}}) = \sigma_I(P_{\text{off}})$), further reducing the number of parameters. Instead, we decided to keep these three parameters free as a control and monitor the relative scale of the $\sigma_I$'s.

Inferring $\Theta$ for each clone individually (i.e., for different genomic separation $s$) is a challenging task, given the large number of parameters (ten parameters) and the risk of overfitting. This is especially true for the clones for which we have limited data ($n \sim 50$ traces for $s = 57$, 82 and 88 kb, see Table S5). However, given our system, most of the parameters could be shared among the clones, since genomic separation is unlikely to affect all the $\sigma_I$'s, $\sigma_R(P)$, and the rates $\{f_2, b_1, b_2, b_3\}$ related to the $P_{\text{off}} \leftrightarrow P_{\text{on}}$ transitions and the homie-homie unbinding transition $P_{\text{off}} \rightarrow O_{\text{off}}$. To drastically simplify our inference problem, we thus initially made the assumption that only $f_1$ and $\sigma_R(O_{\text{off}})$ are clone-specific (and thus

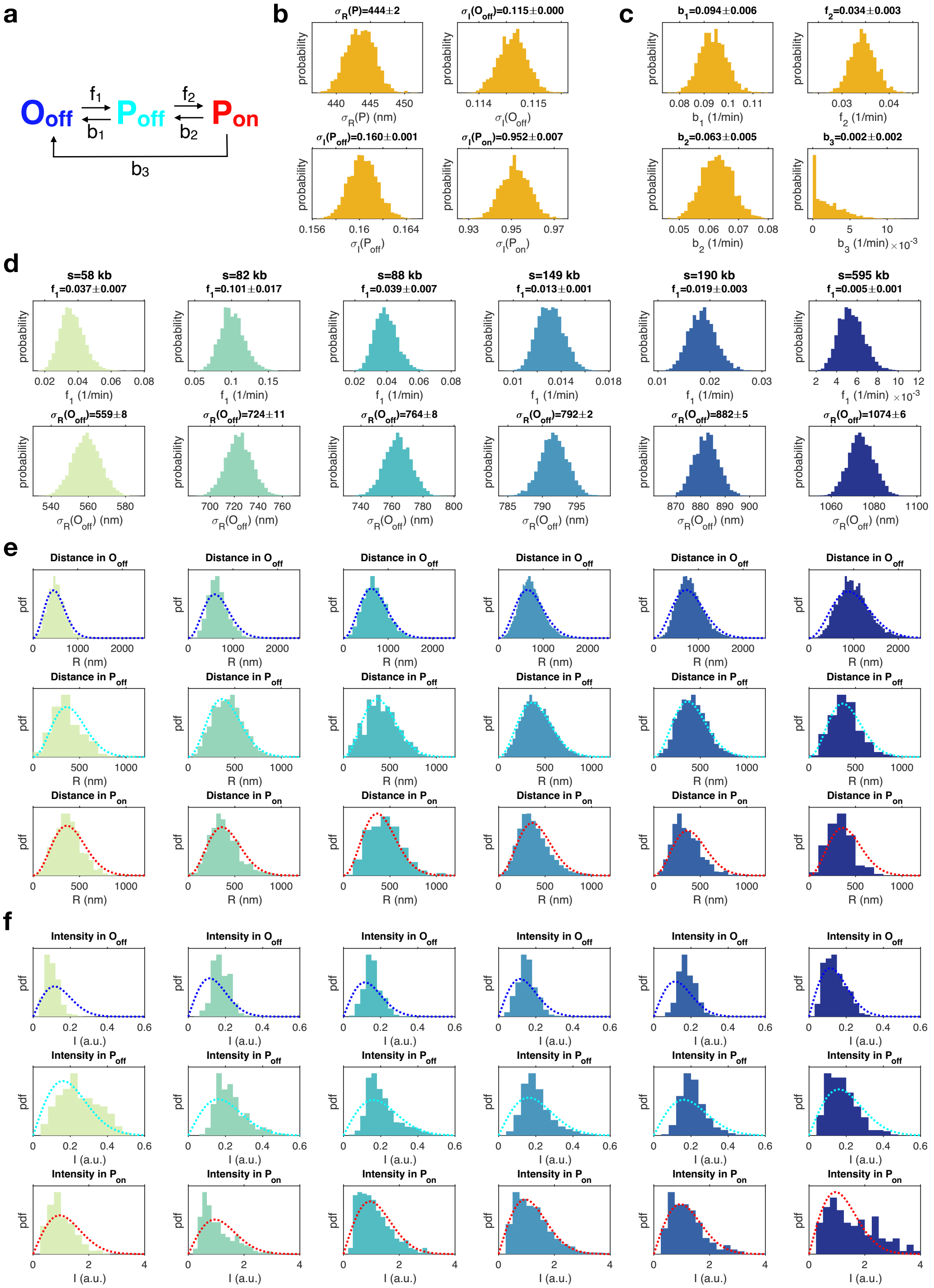

Figure S5: **Calibration of a hierarchical HMM model to infer topological and transcriptional states.** **a**, Schematic of the latent state transitions learned. **b**, Posterior distributions $P(\Theta|D)$ of the global parameters $\Theta = \{\sigma_R(P), \sigma_I(O_{\text{off}}), \sigma_I(P_{\text{off}}), \sigma_I(P_{\text{on}})\}$ related to the observation distributions. **c**, Posterior distributions $P(\Theta|D)$ of the transition rates that are shared among the different clones, i.e., parameters that are assumed independent of genomic separation. **d**, Posterior distributions $P(\Theta|D)$ of the parameters that are specific to each clone, i.e., the distance $\sigma_R(O_{\text{off}})$ and the rate $f_1$. **e**, Comparison between the calibrated observation distributions (dotted lines, Eq. (S4) using the inferred $\sigma_R(O_{\text{off}})$ and $\sigma_R(P)$) and the underlying state based empirical distributions for distances. **f**, Comparison between the calibrated observation distributions (dotted lines, Eq. (S5) using the inferred $\sigma_I(O_{\text{off}})$, $\sigma_I(P_{\text{off}})$ and $\sigma_I(P_{\text{on}})$) and the underlying state based empirical distributions for intensities.

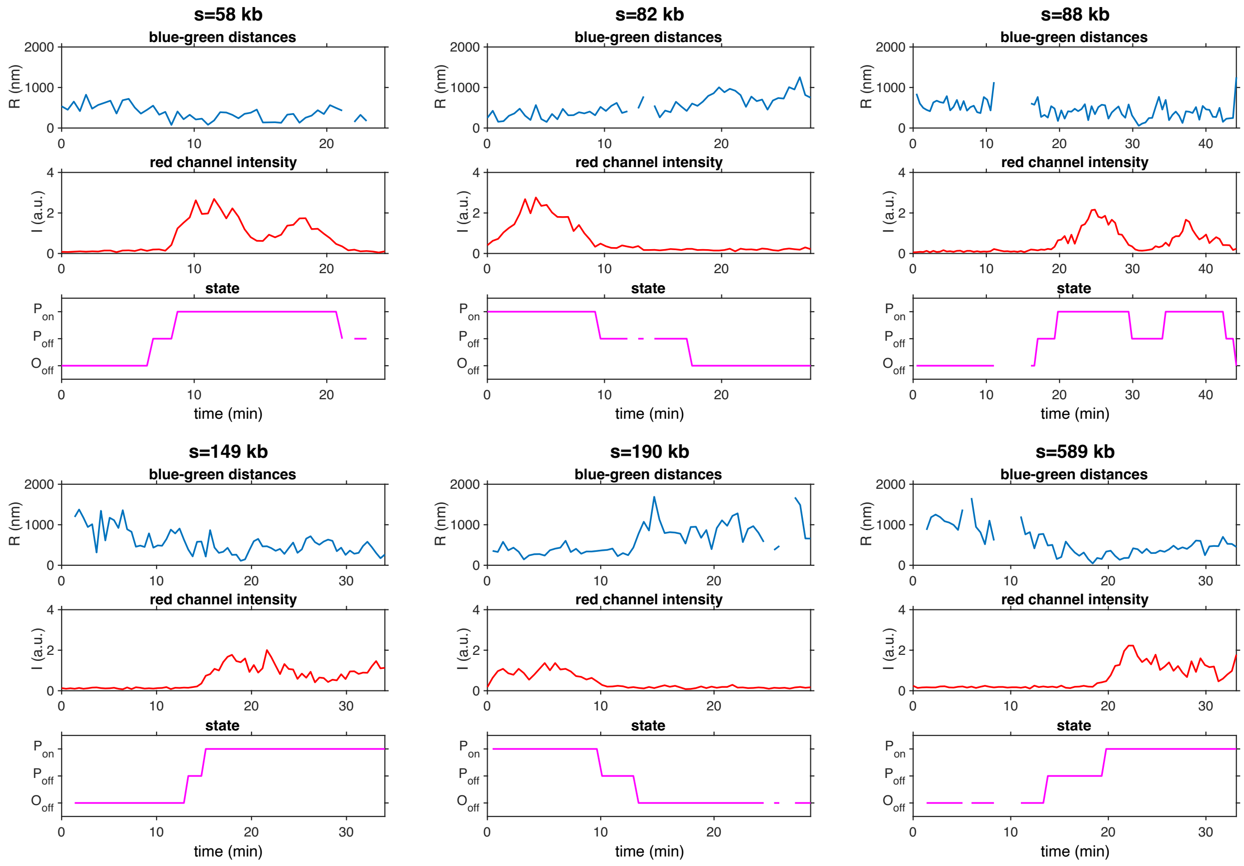

Figure S6: **Example trajectories for various genomic separations.** For each genomic separation $s$, one input pair of blue-green distance $R$ and red transcriptional fluorescence intensity $I$ are shown on top of the inferred state. The inferred state is obtained as the argmax of the posterior decoding $P(\lambda_t|\mathcal{D})$ computed using the calibrated hierarchical HMM (see Fig. S5).

depend on genomic separation $s$), whereas all the other parameters are shared among the clones. Such a structure defines a hierarchical model characterized by the following likelihood function:

$$P(\mathcal{D}|\Theta_H) = \prod_{j=1}^{N_s} P(\mathcal{D}(s_j)|\Theta = \{\Theta_G, \Theta_j\}), \tag{S8}$$

where $\mathcal{D} = \{\mathcal{D}(s_1), ..., \mathcal{D}(s_{N_s})\}$ is the full data set made of all the genomic separation $s_j$, $P(\mathcal{D}(s_j)|\Theta)$ is the likelihood of the data for a specific $s_j$ (see Eq. S2), $\Theta_H = \{\Theta_G, \Theta_1, ..., \Theta_{N_s}\}$ is the ensemble of parameters, among which $\Theta_G = \{\sigma_R(P), \sigma_I(O_{\text{off}}), \sigma_I(P_{\text{off}}), \sigma_I(P_{\text{on}}), f_2, b_1, b_2, b_3\}$ are the global parameters that are shared for all $s_j$, and $\Theta_j = \{\sigma_R(O_{\text{off}}), f_1\}$ are the local parameters specific to each $s_j$. Notably, this assumed hierarchical structure leads to a massive reduction in number of parameters, namely $\#\Theta = 8 + 2 \cdot N_s$ instead of $\#\Theta = 10 \cdot N_s$ (with $N_s = 6$, $\#\Theta = 20$ versus $\#\Theta = 60$ respectively).

We first estimated the joint posterior distribution $P(\Theta_H|\mathcal{D})$ of our hierarchical model (S8). The posterior $P(\Theta_H|\mathcal{D})$ was sampled using a MCMC sampler [9], which enables estimation of the marginal posterior distribution for each parameter. All the marginal posterior distributions are shown in Fig. S5b-d, and the resulting parameter estimates (mean and standard deviation) can be found in Table S2 and S3. Interestingly, the posterior of the rate $b_3$ looks like the prior (log-unfiorm) and the mean rate $b_3$ is much smaller than $b_2$. This clearly indicates that the contribution of $b_3$ to the model dynamics is negligible and

| $\sigma_R(P)$ (nm) | $\sigma_I(O_{\text{off}})$ (a.u.) | $\sigma_I(P_{\text{off}})$ (a.u.) | $\sigma_I(P_{\text{on}})$ (a.u.) |
|---|---|---|---|
| $444 \pm 2$ | $0.115 \pm 0.000$ | $0.160 \pm 0.001$ | $0.952 \pm 0.007$ |
| $b_1$ (1/min) | $f_2$ (1/min) | $b_2$ (1/min) | $b_3$ (1/min) |
| $0.094 \pm 0.006$ | $0.034 \pm 0.003$ | $0.063 \pm 0.005$ | $0.002 \pm 0.002$ |

Table S2: **Global parameters $\Theta_G$ of the hierarchical HMM model estimated using MCMC.**

| $s$ (kb) | $\sigma_R(O_{\text{off}})$ (nm) | $f_1$ (1/min) |
|---|---|---|
| 58 | $559 \pm 8$ | $0.037 \pm 0.007$ |
| 82 | $724 \pm 11$ | $0.101 \pm 0.017$ |
| 88 | $764 \pm 8$ | $0.039 \pm 0.007$ |
| 149 | $792 \pm 2$ | $0.013 \pm 0.001$ |
| 190 | $882 \pm 5$ | $0.019 \pm 0.003$ |
| 595 | $1074 \pm 6$ | $0.005 \pm 0.001$ |

Table S3: **Local parameters $\Theta_j$ of the hierarchical HMM model estimated using MCMC.**

$b_3$ can safely considered to be zero. Moreover, we found that $\sigma_I(O_{\text{off}}) \simeq \sigma_I(P_{\text{off}}) < \sigma_I(P_{\text{on}})$ as expected.

Leveraging the inferred parameters of the hierarchical model, we then reconstructed the states underlying the inter-locus distance and transcriptional activity of individual nulcei using a Viterbi algorithm and posterior decoding [8]. These algorithms take as input the time series of the distances $R$ and red fluorescence values $I$ of a single nucleus. The Viterbi algorithm finds a single state trajectory $\Lambda = (\lambda_0, \lambda_1, ...)$ that maximizes $P(\Lambda|\mathcal{D}_i, \Theta)$, namely the most likely trajectory given the data and the inferred parameters. Alternatively, we also used posterior decoding that enables computation of the state marginal distribution $P(\lambda_t|\mathcal{D}_i, \Theta)$ at time $t$ given the data and the inferred parameters. A single state trajectory $\Lambda = (\lambda_0, \lambda_1, ...)$ can then be computed from the state marginal distribution as $\lambda_t = \text{argmax}\, P(\lambda_t|\mathcal{D}_i, \Theta)$. It turns out that for our system, both approaches, either the Viterbi or Posteriro deconding, predict essentially the same underlying latent state trajectories (see example in Fig. 2c,d and Fig. S6).

Using the state inference, we then checked whether the emission probabilities $P(R|\lambda)$ and $P(I|\lambda)$ (Eq. S4 and S5) calibrated with the inferred parameters $\{\sigma_R(O_{\text{off}}), \sigma_R(P)\}$ and $\{\sigma_I(O_{\text{off}}), \sigma_I(P_{\text{off}}), \sigma_I(P_{\text{on}})\}$ match the empirical distribution (Fig. S5e,f). Overall, the agreement is excellent supporting the validity of our inference. Lastly, to test our assumption behind our hierarchical model, that most parameters

| $s$ (kb) | $\sigma_R(O_{\text{off}})$ (nm) | $\sigma_R(P)$ (nm) | $f_1$ (1/min) | $b_1$ (1/min) | $f_2$ (1/min) | $b_2$ (1/min) |
|---|---|---|---|---|---|---|
| 58 | $560 \pm 7$ | $404 \pm 6$ | $0.023 \pm 0.007$ | $0.079 \pm 0.023$ | $0.110 \pm 0.026$ | $0.063 \pm 0.014$ |
| 82 | $738 \pm 13$ | $464 \pm 5$ | $0.089 \pm 0.019$ | $0.058 \pm 0.011$ | $0.032 \pm 0.006$ | $0.098 \pm 0.019$ |
| 88 | $770 \pm 8$ | $476 \pm 7$ | $0.044 \pm 0.009$ | $0.099 \pm 0.020$ | $0.035 \pm 0.009$ | $0.075 \pm 0.021$ |
| 149 | $793 \pm 2$ | $440 \pm 3$ | $0.014 \pm 0.001$ | $0.098 \pm 0.008$ | $0.029 \pm 0.003$ | $0.066 \pm 0.007$ |
| 190 | $876 \pm 5$ | $420 \pm 6$ | $0.014 \pm 0.003$ | $0.089 \pm 0.016$ | $0.036 \pm 0.008$ | $0.046 \pm 0.009$ |
| 595 | $1071 \pm 6$ | $378 \pm 10$ | $0.005 \pm 0.001$ | $0.124 \pm 0.032$ | $0.020 \pm 0.011$ | $0.050 \pm 0.025$ |

Table S4: **HMM parameters $\Theta$ estimated separately for each genomic separation $s$ using MCMC.**

are shared, we perform inference for each genomic separation separately. To do so, we fixed $\sigma_I(O_{\text{off}})$, $\sigma_I(P_{\text{off}})$ and $\sigma_I(P_{\text{on}})$ to the values of the calibrated hierarchical model (Table S2) and we set $b_3 = 0$ as this parameter appeared unnecessary (see paragraph above). We then re-estimated $f_1$, $f_2$, $b_1$, $b_2$, $\sigma_R(O_{\text{off}})$ and $\sigma_R(O_P)$ for each clone individually and these can be found in Table S4. Overall, the resulting parameter values are very close to the estimates of the hierarchical model (see also section 2.5, and Fig. S10), thus supporting our key assumption.

#### 2.2 Validating the HMM using simulated data

To validate our reconstruction of the states and the inference of the underlying transition rates, we tested our HMM approach on simulated data. We aimed to generate synthetic data sets that mimic real data, namely simulated data that satisfy the same latent state transition with realistic rates $\{f_1, b_1, f_2, b_2, b_3\}$ and observation parameters $\{\sigma_R(O_{\text{off}}), \sigma_R(P), \sigma_I(O_{\text{off}}), \sigma_I(P_{\text{off}}), \sigma_I(P_{\text{on}})\}$ (Fig. S5).

We chose to test our approach on three different genomic separations; short ($s = 58$ kb), medium ($s = 149$ kb) and long ($s = 595$ kb). For each $s$, we generated 300 time-series of blue-green distance $R$ and transcriptional intensity $I$ , whose duration equalled 40 min and sampling time $\Delta t \simeq 30$ sec as in real data. Time-series for the blue-green distance were generated using simulations of a Rouse polymer model with 1000 beads. We tracked the position of two beads equidistant from the central bead of the polymer, with a separation $s/2$ from the center such that the "genomic distance" was given by $s = 58, 149, 595$ beads. We then rescaled time and length-scales such that the relaxation time of the polymer $\tau = s^2$ and the average inter-locus distance matched the experimental values for each value of $s$. To capture the dependence of the transition from the $O_{\text{off}}$ to the $P_{\text{off}}$ state on the dynamics of the 3D distance, we defined a simple implementation of distance-dependent homie-homie binding. Specifically, we defined a success rate of homie-homie binding $r$, such that the two ends of the polymer bind with probability $r$ when in close proximity (defined by a threshold on the 3D distance of 400 nm). The parameter $r$ was assumed constant for all simulations and set to 0.1% to approximately reproduce the $f_1$ rate estimated from real data. Note that this parameter $r$ cannot be compared to the biological success rate of real homie binding, as it depends on the time-scale at which the system is sampled. Once the two homie elements are bound, we include an additional linear spring linking the two loci, with spring constant $k_{\text{link}} = 0.02k$, where $k = 1$ is the spring constant of the backbone of the polymer. The parameter $k_{\text{link}}$ was chosen to generate distances $R$ whose mean $\langle R \rangle \simeq 400$ nm in the $P_{\text{off}}$ and $P_{\text{on}}$ state. In addition, once the $P_{\text{off}}$ state is reached, we ran a Gillespie algorithm to simulate the transition from the $P_{\text{off}}$ to $P_{\text{on}}$ state and from the $P_{\text{off}}$ back to the $O_{\text{off}}$ state [12]. The Gillespie simulation was performed according to the state transition in Fig. S5a, with rates set by the global parameters determined from real data (Table S2 and $b_3 = 0$).

Next, to simulate the intensities $I$ in the $O_{\text{off}}$ and $P_{\text{off}}$ state (i.e., the background), we generate uncorrelated samples from a Rayleigh distribution with parameter $\sigma_I(O_{\text{off}})$ and $\sigma_I(P_{\text{off}})$ set from Table S2, respectively. Lastly, to simulate the transcriptional intensities in the $P_{\text{on}}$ state, we sampled in log-space from a Gaussian process $\tilde{I}(t)$ with mean $\tilde{\mu}$, variance $\tilde{\sigma}^2$ and covariance function $K(t, t')$. We chose $K(t, t') = \exp(-|t - t'|/\tau_e)$ (Ornstein?Uhlenbeck) and we set the correlation time to $\tau_e = 2$ min, which approximates the temporal correlation in the transcriptional signal. The mean $\tilde{\mu}$ and the variance $\tilde{\sigma}^2$ were defined such that the intensity time-series in real-space $I(t) = \exp(\tilde{I}(t))$ is log-normal distributed with mean and variance equal to a Rayleigh distribution of parameter $\sigma_I(P_{\text{on}})$ determined from Table

S2. Namely, $\tilde{\mu}$ and $\tilde{\sigma}^2$ are given by:

$$\tilde{\mu} = \log\left(\frac{\pi\sigma_I(P_{\text{on}})}{2\sqrt{2}}\right) \quad \text{and} \quad \tilde{\sigma}^2 = \log\left(1 + \frac{4-\pi}{\pi}\right) \tag{S9}$$

All the simulated time-series were assumed to start in the $O_{\text{off}}$ state. As an example, a pair of simulated distance $R$ and intensity $I$ time series is shown in Fig. S7c.

With these three simulated data sets in hand ($s = 58, 149$ and $s = 595$ kb), we first aimed to compare the inferred parameters to the input ones used to generate the data. We performed the HMM inference for each data set separately, by sampling the joint posterior distribution $P(\Theta|\mathcal{D})$ with MCMC as described in section 2.1. Thus, for each data set, we estimated the ten free parameters all at once, namely $\Theta = \{\sigma_R(O_{\text{off}}), \sigma_R(P), \sigma_I(O_{\text{off}}), \sigma_I(P_{\text{off}}), \sigma_I(P_{\text{on}}) f_1, f_2, b_1, b_2, b_3\}$. The resulting marginal posterior distributions for $s = 149$ kb are displayed in Fig. S7a with the input (true) parameters highlighted by a vertical dashed line. For each simulated data sets, we estimated the relative error on all parameters. Overall, the absolute relative error for the rates are 30% on average (Fig. S7b top), while for the observation parameters these are 7% on average (Fig. S7b bottom). Thus, even when estimating all the parameters at once (hardest case), the errors remain reasonable and clearly allow us to distinguish different scales among the parameters.

Having checked our ability to infer parameters correctly, we then tested whether our state inference was accurate. We thus performed posterior decoding using our trained HMM on the three simulated data sets, as described in section 2.1. The whole state reconstruction for $s = 149$ kb is shown in Fig. S7d,e; the inferred state trajectories are visually hardly distinguishable from the ground truth. Assessing our reconstruction error on all simulated data sets, we recovered the correct state for more than 95% of the time points (Fig. S7f). Overall, the state inference is excellent and does not strongly depend on the underlying error on the rate parameters, since the signatures of the different states in the data are rather strong (especially for $O_{\text{off}}$ and $P_{\text{on}}$).

##### 2.3 Single trajectory state analysis

Here, we demonstrate that the majority of single-cell trajectories explore the full range of interlocus distances that are sampled by the whole ensemble of trajectories in all three inferred states. From the ensemble of state trajectories (Fig. S8a), we plot the distance distributions for each single trajectory as a function of its inferred transcriptional and topological state (Fig. S8b-c). We find that in the majority of cases, the single trajectory distributions approximately match the population averaged distribution, suggesting that the single-cell trajectories explore the full range of distances of each state.

As a more stringent criterion, we perform a hypothesis test of the null hypothesis that the single-cell distribution is indistinguishable from the ensemble averaged distribution. Specifically, we use the two-sided Kolmogorov-Smirnov test comparing for each trajectory the single-trajectory distribution to the population distribution. The null-hypothesis that the distributions are the same is rejected if $p < 0.05$. We find that for most data sets, for the majority of trajectories in all states for all genomic separations, the null-hypothesis is not rejected (Fig. S8e).

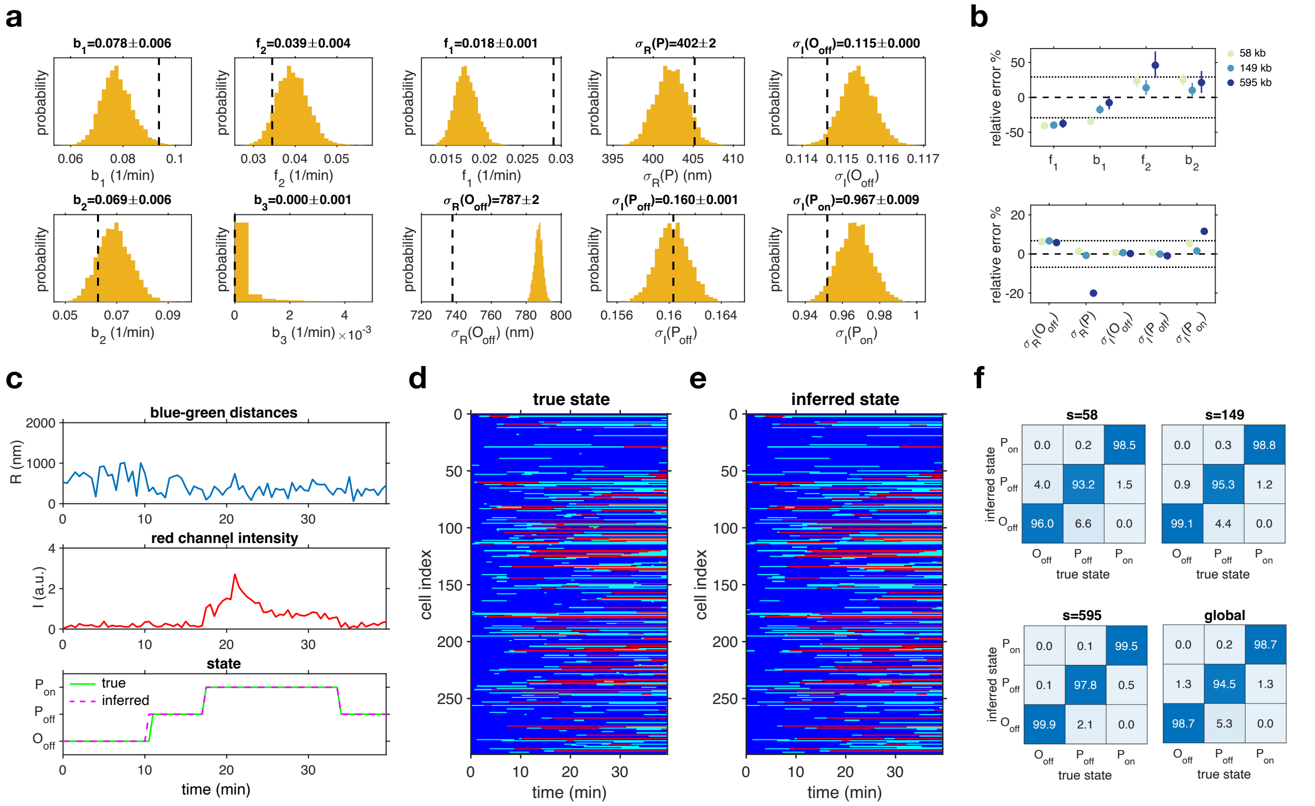

Figure S7: **Validating the HMM inference on simulated data.** **a**, Posterior distributions $P(\Theta|D)$ estimated using simulated data with genomic separation $s = 149$kb. Vertical dashed lines indicate the input parameters used for generating the data. **b**, Relative error (in %) with respect to input values on the inferred rates (top) and distances (bottom) for three simulated data sets with different genomic separation (color code; $s = 58$, 149 and 595kb). **c**, State inference on simulated data. **d**, True states for the entire simulated data set $s = 149$kb. **e**, Inferred states for the entire simulated data set $s = 149$kb. **f**, Fraction of inferred states as a function of true input (in %) for all simulated data sets.

#### 2.4 Estimation of transcriptional lifetime

To estimate the lifetime of transcriptional activity, we use the inferred state trajectories (section 2.1) to calculate the typical time the system remains in the $P_{\mathrm{on}}$ state. Lifetime estimation is challenging due to the problem of censoring, referring to the fact that trajectories may start or end in the $P_{\mathrm{on}}$ state, which gives only partial access to the lifetime in such a trajectory. To deal with this problem, we use the Kaplan-Meier estimator for the survival function of the transcription lifetimes [13]. The survival function $S(t)$ gives the probability that the transcriptionally active state has *not* transitioned to a transcriptionally inactive state, and is defined as

$$S(t) = 1 - \int_0^t p(t')dt' \tag{S10}$$

where $p(t)$ is the probability distribution of transcriptional lifetimes. We use the Kaplan-Meier estimator [13, 14]

$$\hat{S}(t) = \prod_{t_i<t} \left(1 - \frac{N^{(i)}_{\mathrm{uncensored}}}{N^{(i)}}\right) \tag{S11}$$

where $N^{(i)}_{\mathrm{uncensored}}$ is the number of uncensored events of length $t_i$ and $N^{(i)}$ is the number of events with length larger than $t_i$. From this estimator, we obtain the survival curves shown in main text Fig. 2c

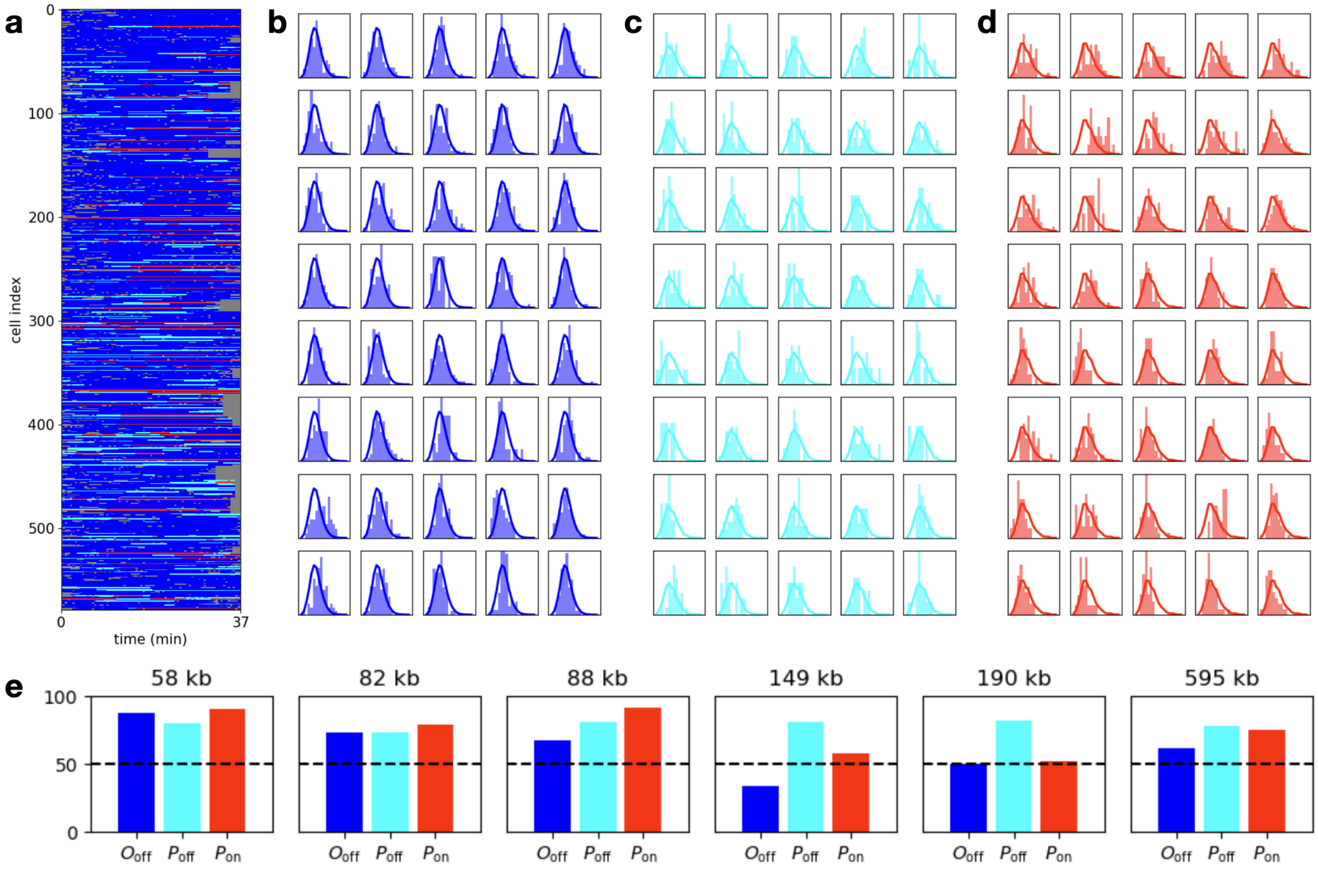

Figure S8: **Single-trajectory interlocus distance distributions.** **a**, All state trajectories for genomic separation $s = 149$ kb. **b-d**, Single trajectory interlocus distance distributions for the three states ($s = 149$ kb). Each panel shows the distance distribution of a single trajectory as a bar histogram, and the population averaged distribution as a solid line. Only trajectories with at least 10 time-points are included to ensure sufficient statistics for comparison. **e**, Percentage of trajectories for which the null-hypothesis that the single-cell and the population distribution are the same is not rejected, for each state and genomic separation.

and Fig. S9a. From these curves, the estimate of the median lifetime can be determined by finding the intercept where $\hat{S}(\text{median}[\tau_{\text{trans}}]) = 0.5$.

As a second method to estimate the median lifetime is a maximum likelihood estimate under the assumption that the lifetime distribution is exponential, i.e.

$$p(t) = \frac{1}{\langle \tau_{\text{trans}} \rangle} e^{-t/\langle \tau_{\text{trans}} \rangle} \tag{S12}$$

where $\langle \tau_{\text{trans}} \rangle$ is the mean transcriptional lifetime. Then, the maximum likelihood estimate of $\langle \tau_{\text{trans}} \rangle$ is given by [14]

$$\widehat{\langle \tau_{\text{trans}} \rangle} = \frac{1}{N_{\text{uncensored}}} \sum_i t_i \tag{S13}$$

and the median lifetime is given by $\ln(2)\widehat{\langle \tau_{\text{trans}} \rangle}$.

The median lifetime estimates from both methods are given in main text Fig. 2d. For the genomic separation of 595 kb, there are only 5 observed $P_{\text{on}}$ states out of a total of 164 trajectories, meaning that the lifetime is hard to estimate for this sample (see last panel of Fig. S9a). For 3.3 Mb, there are no observed $P_{\text{on}}$ states.

Furthermore, we compare these estimates to simply taking the mean of all observed lifetimes (Fig. S9b), showing that also in this case, there is no variation with genomic separation, and the typical lifetime is $\approx$ 10 min.

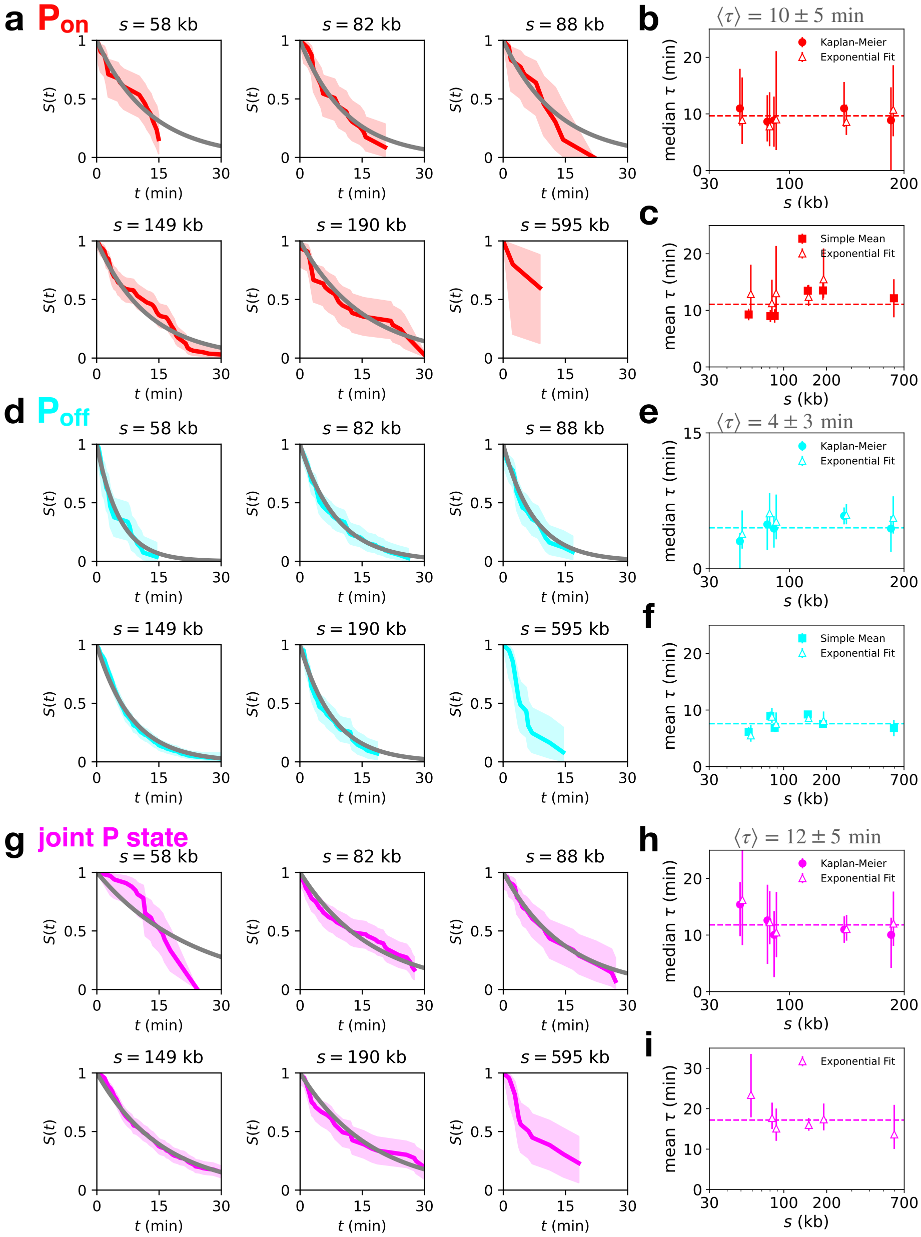

Figure S9: **Lifetime-estimation using survival probability curves.** **a**, Kaplan-Meier survival curves for each genomic separation. Shaded regions: 95% confidence intervals [14]. Grey lines: exponential fits obtained by using the maximum likelihood mean lifetime (using Eq (S13) in Eq. (S12)). **b**, Median lifetimes obtained by the exponential fit (triangles), or by reading off the time at which the Kaplan-Meier survival curves cross $S = 0.5$ (dots). Equivalent to main text Fig. 2d for the $P_{\text{on}}$ state. The mean of median lifetimes across genomic separations is noted above (error bars: standard deviation calculated from total variance). **c**, Mean lifetimes obtained by the exponential fit (triangles) and by simply taking the mean of all observed lifetimes (squares). Errorbars on exponential fit results represent 95% confidence intervals [14], and on means represent bootstrap error bars. Panels **a-c**, are for the $P_{\text{on}}$ state (red), while panels **d-f** show the same plots for the $P_{\text{off}}$ state (cyan), and panels **g-i** for the joined paired state when the system is either in $P_{\text{on}}$ or $P_{\text{off}}$ (magenta).

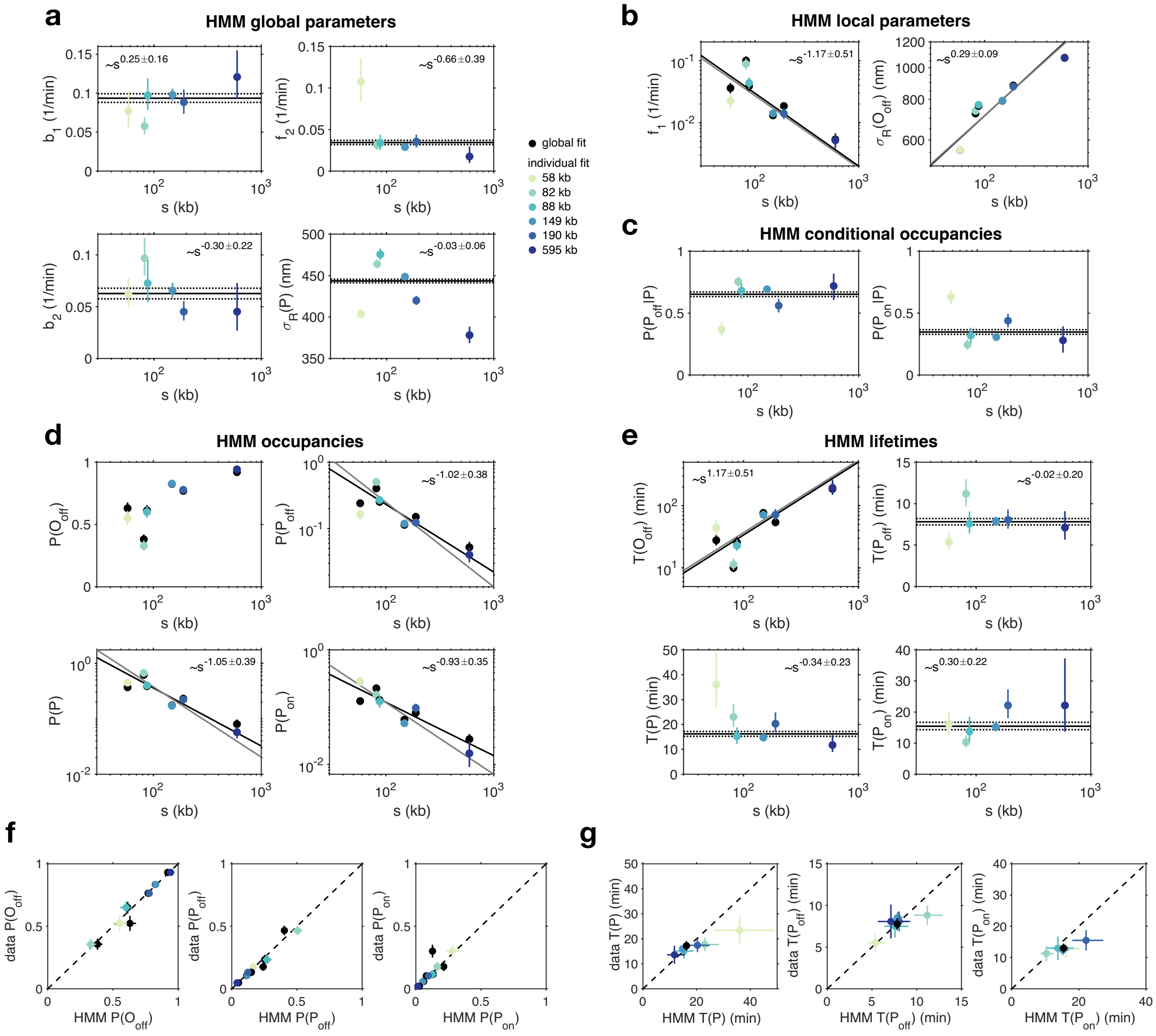

Figure S10: **Scaling of the HMM parameters, the predicted state-occupancy and state-lifetime. a-b**, Inferred parameters as a function of genomic separation $s$. **a**, These parameters were assumed shared among the clones in the hierarchical model. Performing the inference separately for each individual clone lead to the parameter values that remain close to the ones obtained using the hierarchical model (black solid and dashed line, mean and standard deviation respectively). **b**, These parameters were assumed to scale with genomic separation $s$. Performing the inference separately for each individual clone preserve the scaling observe with the hierarchical model (gray line, power-law fit). **c**, HMM-predicted conditional occupancy of $P_{\text{off}}$ and $P_{\text{on}}$ states as a function of genomic separation $s$. **d**, HMM-predicted occupancy of the different states as a function of genomic separation $s$. **e**, HMM-predicted residence time within the different states as a function of genomic separation $s$. **f**, Comparison between HMM-predicted and measured occupancies. **g**, Comparison between HMM-predicted and measured lifetimes.

#### 2.5 HMM parameter scaling analysis

To assess which parameters of our HMM model scale with genomic separation $s$, we systematically compare the parameters of our global inference using the hierarchical model to the parameters obtained by doing inference on each $s$ separately (see also section 2.1). The hierarchical model was built on the assumption that most parameters are independent of $s$, and thus can be shared across clones (Eq. S8).

We first analyzed the parameters that were assumed to not scale with $s$ (Fig. S10a), namely the distance

$\sigma_R(P)$ and the rates $\{f_2, b_1, b_2\}$ related to the $P_{\text{off}} \leftrightarrow P_{\text{on}}$ transitions and the homie-homie unbinding transition $P_{\text{off}} \rightarrow O_{\text{off}}$. Overall, these parameters, when inferred separately for each $s$, remain numerically very close to the global estimate (black line) made with the hierarchical model. In addition, the power-law exponents characterizing the scaling of these parameters as function of $s$ are close to zero (within error bars), validating our assumption that these are mostly independent of $s$. Next, we investigated the parameters that were assumed to scale with $s$ (Fig. S10b), namely the open distance $\sigma_R(O_{\text{off}})$ and the rate $f_1$. When these parameters are inferred separately for each $s$, we again recover values that are numerically very close to the global estimate (black dots). The observed scaling of the parameters with $s$ are very similar between the two inference approaches (hierarchical vs individual model, black and gray lines respectively). Moreover, the estimated scaling exponents are significantly different from zero and match the estimation based on data (Fig. 2).

Next, we investigated the impact of the parameter scaling on the steady state occupancy (i.e., the probability) of the three states. To calculate the occupancy, we solved analytically the steady state equation $GP = 0$, where $G$ is the Laplacian matrix of the system (Eq. S6). We obtained the following expression for the probabilities as function of the rates:

$$P(O_{\text{off}}) = \frac{b_1 b_3 + b_1 b_2 + f_2 b_3}{Z} \tag{S14}$$

$$P(P_{\text{off}}) = \frac{(b_3 + b_2) f_1}{Z} \tag{S15}$$

$$P(P_{\text{on}}) = \frac{f_2 f_1}{Z} \tag{S16}$$

$$Z = b_1 b_3 + b_1 b_2 + f_2 b_3 + (b_3 + b_2 + f_2) f_1. \tag{S17}$$

Using the equations above, the conditional probabilities $P(P_{\text{off}}|P)$ and $P(P_{\text{on}}|P)$ are easily calculated:

$$P(P_{\text{off}}|P) = \frac{b_3 + b_2}{f_2 + b_3 + b_2} \tag{S18}$$

$$P(P_{\text{on}}|P) = \frac{f_2}{f_2 + b_3 + b_2} \tag{S19}$$

(S20)

Interestingly, both $P(P_{\text{off}}|P)$ and $P(P_{\text{on}}|P)$ do not depend on $f_1$, and thus are not expected to vary with genomic separation $s$. Indeed, when computing these conditional probabilities using the parameters of the hierarchical model (Fig. S10c, black line), we found $P(P_{\text{off}}|P) = 0.65$ and $P(P_{\text{on}}|P) = 0.35$, which are independent on $s$ and match the values estimated from data Fig. 2e. Similarly, when computing these probabilities using the parameters from the individual fits (Fig. S10c, color dots), the probabilities cluster around the global estimate (albeit $s = 58$ kb appears to be an outlier).

We then investigated the scaling of the occupancies themselves (Eq. S14). Given our parameter regime, we expect the following scaling for the probabilities: $P(O_{\text{off}}) \sim 1/(c + f_1)$ and $P(P) \sim P(P_{\text{off}}) \sim P(p_{\text{on}}) \sim f_1$, where $f_1 \sim s^{-1.17 \pm 0.51}$ (see Fig. S10b). We computed these probabilities using both the hierarchical and individual fit parameters and found scaling relationships that are very consistent with our expectation (Fig. S10d).

Next, we investigated the scaling of the lifetime (mean residence time) of the different states. In our HMM framework, the lifetimes are phase-type distributed and are easily computed from the state transition matrix (Eq. S6). We found:

$$T(O_{\text{off}}) = \frac{1}{f_1} \tag{S21}$$

$$T(P_{\text{off}}) = \frac{1}{b_1 + f_2} \tag{S22}$$

$$T(P_{\text{on}}) = \frac{1}{b_2 + b_3} \tag{S23}$$

$$T(P) = \frac{f_2 + b_2 + b_3}{b_1 b_2 + b_1 b_3 + f_2 b_3} \tag{S24}$$

Only $T(O_{\text{off}})$ depends on $f_1$, and thus varies with $s$, whereas $T(P_{\text{off}})$, $T(P_{\text{on}})$ and $T(P)$ are supposed to be mostly independent on $s$. We computed these lifetimes using both the hierarchical and individual fit parameters and found scaling relationships that are again very consistent with our expectation (Fig. S10e). Indeed, $T(P_{\text{off}})$, $T(P_{\text{on}})$ and $T(P)$ are essentially constant, while $T(O_{\text{off}})$ varies as the inverse of $f_1$.

Lastly, we compared the occupancies and lifetimes predicted by the HMM parameters to the occupancies and lifetimes estimated from data based on state splitting (Fig. S10f,g). Overall, the excellent agreement between the different quantities predicted by the HMM parameters and the ones estimated from data (using state splitting) supports the validity of our inference and is indicative of the mathematical consistency of our approach. Importantly, it should be noted that such an agreement is not necessarily trivial. Indeed, the HMM model mainly acts as prior for the state inference, thus if the data possessed features that could not be accounted for by the model, discrepancies would have arisen.

##### 2.6 Results without state splitting

The analysis of the two-locus dynamics shown in the main text is done for the $O_{\text{off}}$ only. To demonstrate that our results on the two-locus dynamics do not sensitively depend on the details of the state inference, we compare the statistics of the $O_{\text{off}}$ state to those obtained from the entire data set comprising all three states. Since the data are dominated by the $O_{\text{off}}$ state, the state splitting does not have a strong impact on most statistics, especially for large genomic separations. As expected, the average distances of the full data are lower than those of the $O_{\text{off}}$ state only, and the inferred two-locus diffusion coefficients $\Gamma_2$ are lower. However, key scaling exponents are not strongly affected, including the scaling of the inferred relaxation times $\tau$ (Fig. S11).

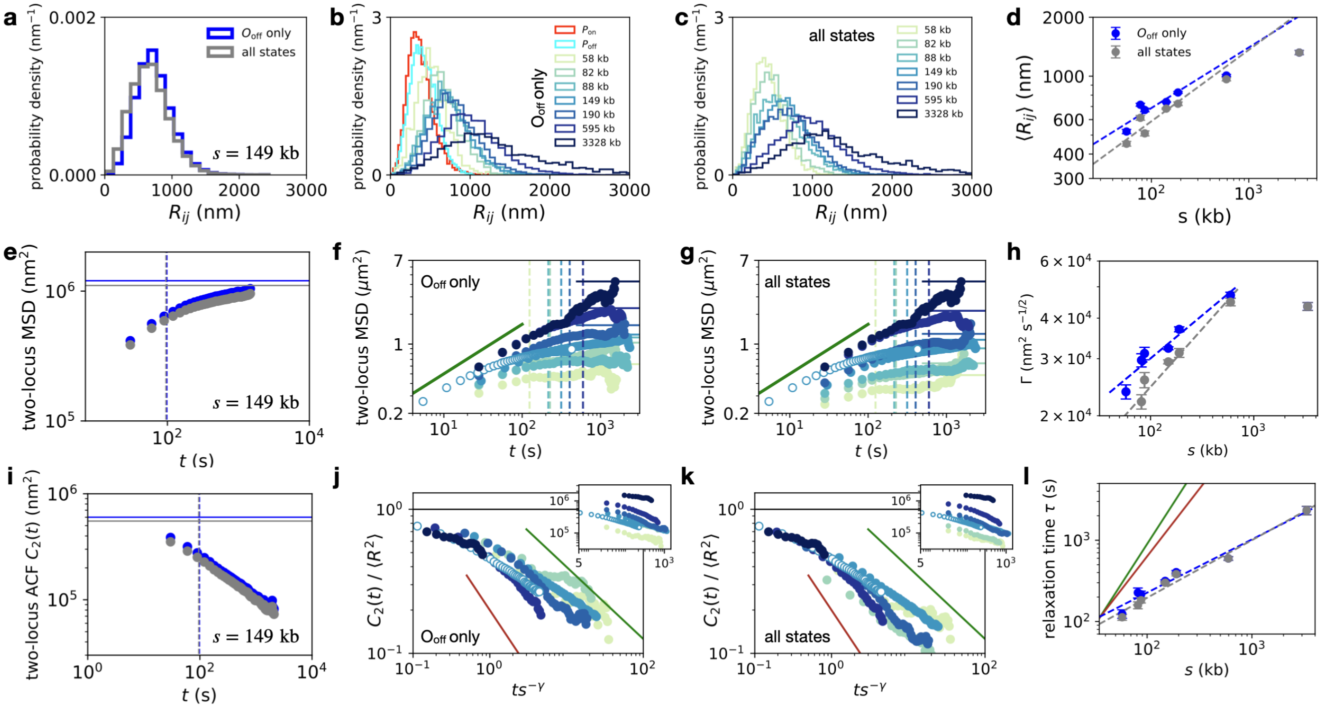

Figure S11: **Two-locus analysis is independent of state inference.** **a**, Probability distributions of the inter-locus distances $R_{ij}$ ($s = 149$ kb). Blue: distribution for the $O_{\text{off}}$ state only, as shown in main text Fig. 2a. Grey: distribution for all states. **b,c**, Probability distributions of the inter-locus distances $R_{ij}$ for all separations, with (b) and without state splitting (c). **d** Scaling of mean interlocus distance with (blue) and without state splitting (grey). **e,f,g**, Two-locus MSD $M_2(t)$, with inferred values of the relaxation time $\tau$ (dashed lines) and the asymptote $2\langle R^2 \rangle$. **h**, Scaling of the two-locus diffusion coefficient with genomic separation with (blue) and without state splitting (grey). **i,j,k**, Two-locus ACF $C_2(t)$, with inferred values of the relaxation time $\tau$ (dashed lines) and the asymptote $2\langle R^2 \rangle$. **h**, Scaling of the inferred relaxation time with genomic separation with (blue) and without state splitting (grey).

#### 2.7 Analysis of no-*homie* data

To test the role of the *homie* insulator-mediated focal contacts for the transcriptional dynamics in our system, we employ a reporter construct in which the *homie* sequence is replaced by a $\lambda$ DNA sequence of the same length. In this system, only two topological states are possible: a transcriptionally inactive configuration $O_{\text{off}}$ and a transcriptionally active configuration $P_{\text{on}}$. To account for this in our state inference, we simplified our HMM hierarchical model by only modeling the $O_{\text{off}} \leftrightarrow P_{\text{on}}$ transitions with rates $f_1$ and $b_1$. Thus, we had only four global parameters $\{\sigma_R(P_{\text{on}}), \sigma_I(O_{\text{off}}P_{\text{on}}), \sigma_I(P_{\text{on}}), b_1\}$ and two local parameters $\{\sigma_R(O_{\text{off}}), f_1\}$ to infer from the no-*homie* data. The resulting inferred parameters are displayed in Fig. S12. The state inference was then performed using this model specifically calibrated on the no-*homie* data.

Based on the state trajectories, we find that there is still frequent transcriptional activity for genomic separation 58 kb ($8.5 \pm 0.8\%$), but almost none ($< 1\%$ of time-points) for 149 kb (Fig. S13a). We furthermore compare the statistics of the 3D interlocus distance between the homie and no-*homie* constructs of the same genomic separation. Due to the different statistics of the state dynamics in the two constructs, we compare data from the $O_{\text{off}}$ state in both cases. We find that distance distributions largely overlap (Fig. S13b). The two-locus MSDs also exhibit a similar shape and are well fitted by the ideal chain theoretical expression, although we observe small, but not systematic, differences in the two-locus diffusion coefficient between homie and no-*homie* constructs (Fig. S13c).

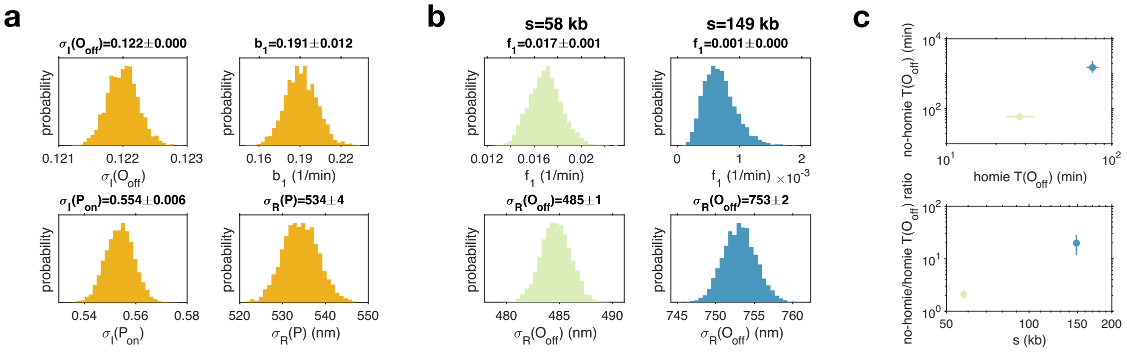

Figure S12: **Calibration of the hierarchical HMM model for the no-*homie* data.** **a**, Posterior distributions of the global parameters $P(\Theta|D)$ estimated using the no-*homie* data with genomic separation $s$ = 58kb and $s$ = 149kb. **b**, Posterior distributions $P(\Theta|D)$ of the clone specific-parameters, which vary with genomic separation $s$. **c**, Comparison of the HMM-predicted open state lifetime $T(O_{\text{off}})$ between the homie and no-*homie* data.

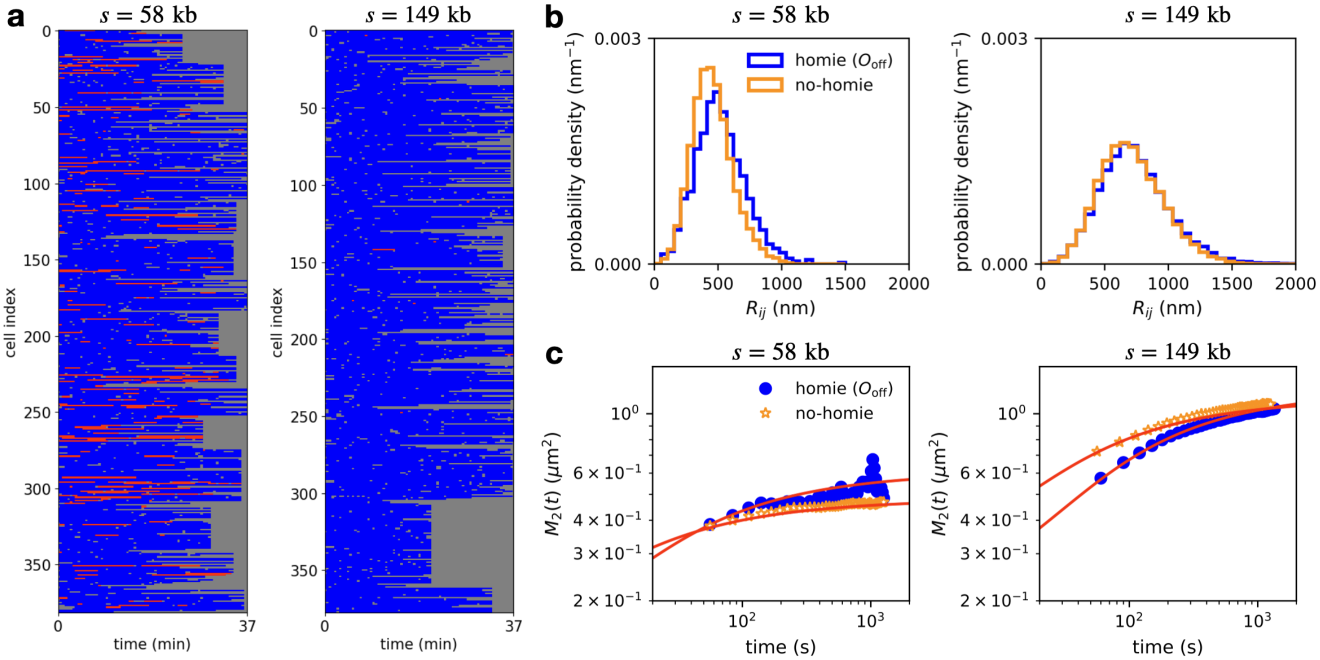

Figure S13: **The open state dynamics is not affected by the presence of homie.** **a**, All state trajectories for no-*homie* constructs with genomic separations 58 and 149 kb. **b**, Probability distributions of the inter-locus distances $R_{ij}$. Blue: distribution for the $O_{\text{off}}$ state of homie constructs, orange: distributions of the transcriptionally inactive state in no-*homie* constructs. **c**, Two-locus MSD $M_2(t)$ with Rouse fits shown as red lines.

### 3 Analysis of single-locus dynamics

The analysis of the motion of single chromosomal loci is only possible if any displacement and rotation of the cell nucleus as a whole is corrected for, as this would affect the apparent dynamics of the DNA loci. This problem does not affect relative measures based on the inter-locus distance vector $\mathbf{R}_{ij}(t)$ (section 4.1). In particular correcting rotations is challenging as at least three loci need to be tracked to infer global rotations. Previous works have therefore focused solely on the statistics of the inter-locus distance [14–16], or, in recent work, relied on multi-locus tracking to correct global movement [17].

In our system, however, the only global mode to be corrected is an overall affine translation of the cell nuclei within the embryo, which we can correct for by nucleus tracking in the $xy$-directions (section 1.3). Nucleus tracking is difficult in the axial $z$-direction. We therefore estimate the statistics of single locus diffusion using the $xy$-components of the dynamics.

To characterize the motion of single loci, we calculate the single-locus mean-square-displacement (MSD) defined in each spatial dimension $i = \{x, y, z\}$ as

$$M_1^{(i)}(t) = \left\langle |r_i(t_0 + t) - r_i(t_0)|^2 \right\rangle_{t_0} = \Gamma_1^{(i)} t^\beta \tag{S25}$$

where averages are averages over time and samples. The 3D MSD is then given by $M_1(t) = M_1^{(x)}(t) + M_1^{(y)}(t) + M_1^{(z)}(t)$. Assuming that all three spatial dimension are identical, we therefore estimate $M_1(t)$ from the $x$ and $y$ components via $M_1(t) = \frac{3}{2}(M_1^{(x)}(t) + M_1^{(y)}(t))$.

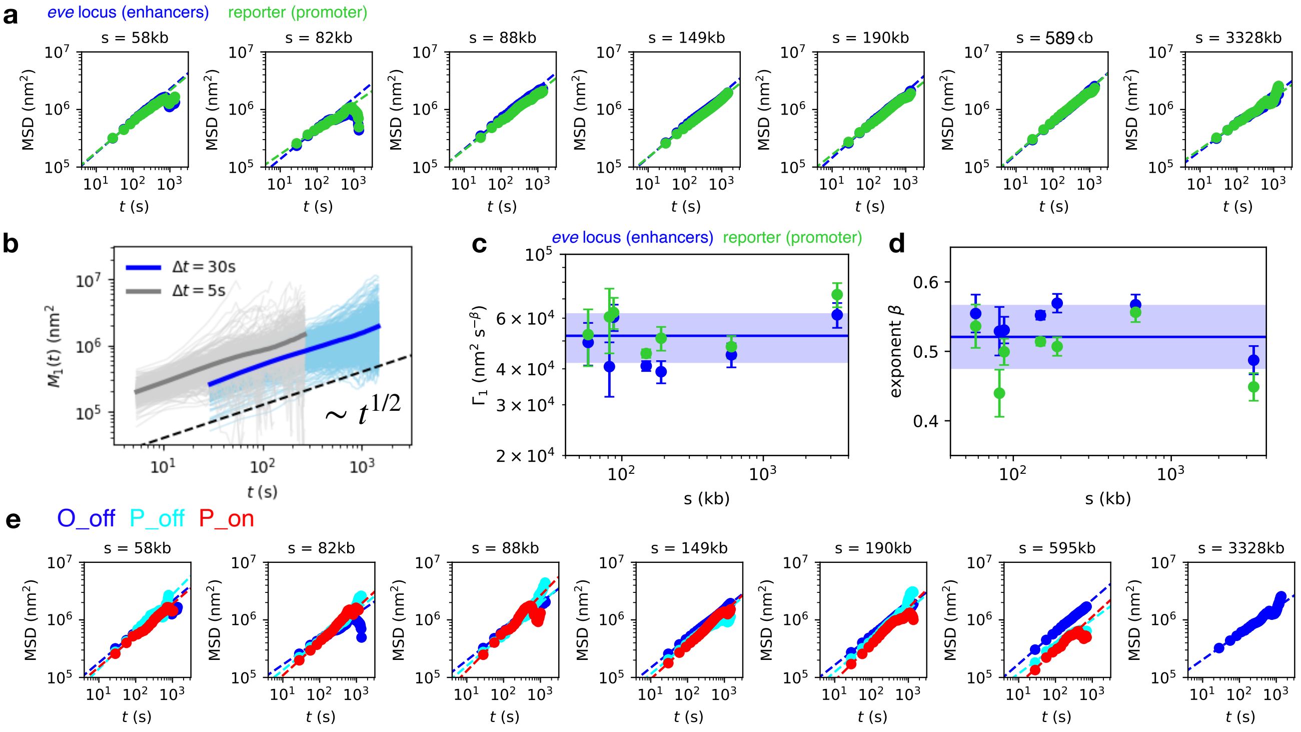

Figure S14: **Analysis of single-locus dynamics.** **a**, Single-locus MSDs $M_1(t)$ for the enhancer (blue) and promoter (green) for each genomic separation $s$, estimated from $xy$-components of the trajectories. Dashed lines indicate linear fits to the first 5 data points. **b**, Single-locus MSDs $M_1(t)$ for $s = 149$kb at two different imaging time-intervals $\Delta t = 5$s (grey) and $\Delta t = 30$s (blue). Thin lines indicate the MSDs of single trajectories. **c**, Single-locus diffusion coefficients $\Gamma_1$ measured using the linear fits in panel a. **d**, Single-locus diffusion exponents $\beta$ measured using the linear fits in panel a. **e**, Single-locus MSDs $M_1(t)$ for the three topological states $O_{\text{off}}$ (blue), $P_{\text{off}}$ (cyan), $P_{\text{on}}$ (red) for each genomic separation $s$. Dashed lines indicate linear fits to the first 5 data points. All MSDs are given in nm$^2$.

We find that the single-locus MSD is consistent across enhancer vs promoter loci, for different genomic separations, for different imaging time-intervals, and for the three inferred topological states (Fig. S14).

Additionally, we also report the velocity autocorrelation functions of single loci [18], given by

$$C_v(t) = \langle \mathbf{v}^{(\delta)}(t_0) \cdot \mathbf{v}^{(\delta t)}(t_0 + t) \rangle_{t_0} \tag{S26}$$

where velocities are calculated over variable time-interval $\delta = n\Delta t$ where $\Delta t$ is the imaging time-interval: $\mathbf{v}^{(\delta)}(t) = (\mathbf{r}(t + \delta) - \mathbf{r}(t))/\delta$. We find that these correlations collapse with $\delta$ and are well described by the Rouse model prediction [18, 19] (Fig. S15):

$$\frac{C_v^{(\delta)}(t)}{C_v^{(\delta)}(0)} = \frac{|t-\delta|^\beta + |t+\delta|^\beta - 2|t|^\beta}{2\delta^\beta} \tag{S27}$$

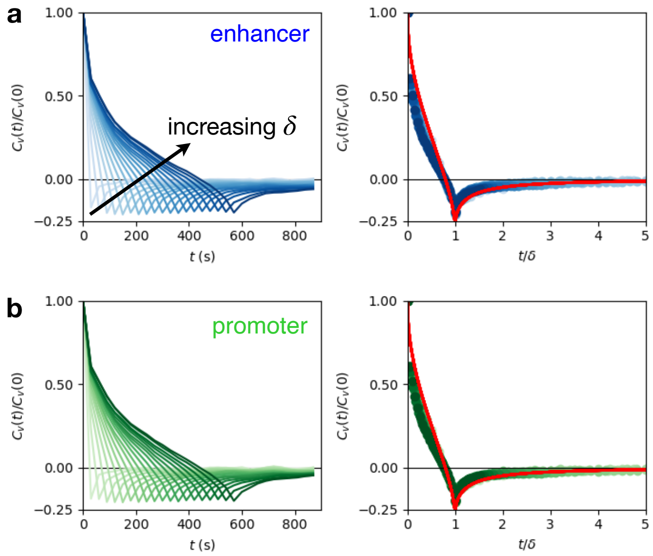

Figure S15: **Velocity autocorrelation functions** $C_v(t)$**.** Left: $C_v^{(\delta)}(t)/C_v^{(\delta)}(0)$ as a function of $t$ with increasing $\delta$. Right: $C_v^{(\delta)}(t)/C_v^{(\delta)}(0)$ as a function of $t/\delta$, showing that the functions collapse. Red line: Theory prediction for the Rouse model (Eq. (S27)) with $\beta = 1/2$. Curves shown here are for $s = 149$ kb. Similar results are obtained for all other genomic separations.

### 4 Analysis of two-locus dynamics

In this section, we provide details of how we test the theoretical predictions from section 5 using experimental data by calculation of two-locus MSDs, inference of the relaxation time (section 4.1) and collapsing the two-locus autocorrelations (section 4.2). Note before any further analysis, we correct $\mathbf{R}_{ij}$ to account for chromatic aberration (section 1.4).

#### 4.1 Two-locus MSD and relaxation time inference

As shown in main text Fig. 3, we find that the early-time scaling of the two-locus MSD exhibits behaviour close to the ideal Rouse exponent $\beta = 1/2$, and the long-time scaling of the two-locus autocorrelations is also close to the ideal Rouse expectation. Thus, we attempt to fit the complete two-locus MSD with the analytical prediction obtained for the ideal chain Rouse model [14]. We use the expression for an infinite continuous polymer; for discussion of the assumption $s \ll N$, see section 5:

$$M_2(t) = 2\Gamma_2 t^{1/2}(1 - e^{-\tau/\pi t}) + 2J\ \mathrm{erfc}[(\tau/\pi t)^{1/2}] \tag{S28}$$

where $J = \langle R^2 \rangle$ and $\tau = (J/\Gamma_2)^2$. Note that we refer to the diffusion coefficient obtained from the two-locus MSD as $\Gamma_2$, while we write $\Gamma_1$ for the single locus diffusion coefficient. For any homogeneous polymer model, such as the ideal or crumpled chains, $\Gamma_1 = \Gamma_2$. However, we do not make this constraint on the data, and indeed find that the fitted $\Gamma_1$ and $\Gamma_2$ differ (Fig. S14).

We find that the two-locus MSDs are well-fitted by Eq. S28, and the fitted value of $J$ coincides with the measured asymptote $\langle R^2 \rangle$ (Fig. S17a). Additionally, the presence of measurement errors in the tracking affects the MSD curves, leading to an additive contribution of $4(\sigma_x^2 + \sigma_y^2 + \sigma_z^2)$, reflecting that the error can be different in each spatial direction. We perform a Bayesian maximum likelihood fit of the two-locus MSD using the python-library tracklib [14] with free parameters $\{J, \Gamma_2, \sigma_z\}$. We use a log-uniform prior for these parameters since they are real positive and have some unknown scale. We do not fit measurement errors in the $x$ and $y$-directions as the estimates for these errors obtained from such a 5-parameter fit are consistent with $\sigma_x = \sigma_y = 0$, and therefore not significant.

To determine the inference errors on the fitted parameters, we perform bootstrapping as described in refs. [20,21]. Briefly, from each data set of $N$ cell trajectories $\{\mathbf{r}_k\}$, where $k = 1...N$, we generate $N_{\mathrm{BS}}$ bootstrap realizations by randomly sampling $N$ trajectories with replacement for each realization. To obtain the error in the parameters $\{J, \Gamma_2, \sigma_z\}$, we infer their value for each bootstrap realization by performing Bayesian MSD fitting on each realization and take the standard deviation of all obtained fit parameters as our estimate for the error in each fit parameter. The distributions of the bootstrap estimates are shown in (Fig. S17b-d), and range between $1 - 6\%$ in the polymer parameters $J, \Gamma_2$, with larger errors in the inferred measurement error amplitude $\sigma_z$. Having obtained our estimate for the relaxation times, we compare them to the time-scales of transitioning from the $\mathrm{O_{off}}$ state to the paired configuration, estimated from the HMM model: $T(\mathrm{O_{off}}) = f_1^{-1}$. We find that these transition times correlate well with the relaxation time (Fig. S16a), and their absolute value is larger by a factor 10 on average (Fig. S16b).

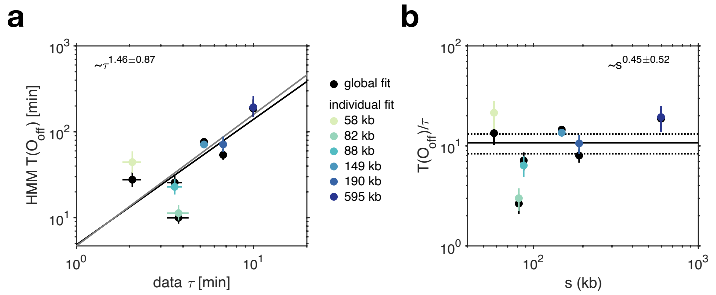

Figure S16: **The relaxation time dictates the open state lifetime.** **a**, Lifetime of the open state $T(O_{\mathrm{off}})$ as a function of the polymer relaxation time $\tau$. $T(O_{\mathrm{off}})$ clearly scales with $\tau$. **b**, Ratio of the open state lifetime $T(O_{\mathrm{off}})$ and the relaxation time $\tau$ as a function of the genomic separation $s$. The ratio is essentially conserved across genomic separation and its value corresponds to a factor 10 on average.

Furthermore, we demonstrate the accuracy of the relaxation time inference using ideal chain Rouse simulations. Our sampling of the two-locus MSD is limited by two key factors: the imaging time interval $\Delta t$ and the maximum trajectory length $t_{\mathrm{max}}$. We aim to infer the relaxation time scaling from a set of genomic separations with minimum and maximum relaxation times $\tau_{\mathrm{min}}$ and $\tau_{\mathrm{max}}$, corresponding to the shortest and longest genomic separation. The inference will be limited if $\Delta t > \tau_{\mathrm{min}}$ or $t_{\mathrm{max}} < \tau_{\mathrm{max}}$. Based on the inference from experimental data, the shortest and longest genomic separation are close to these thresholds (Fig. S17a). To test the accuracy of the inference in this case, we generate simulated data with $t_{\mathrm{max}}/\tau_{\mathrm{max}} = 1$ (as in the experimental data for $s = 3.3$ Mb), and vary $\Delta t/\tau_{\mathrm{min}}$ (Fig. S18a). Experimentally, we have $\Delta t/\hat{\tau}_{\mathrm{min}} = 0.76$ (for $s = 58$ kb). The larger this ratio, the more difficult the inference. However, we find that the inference performs well for ratios as high as $\Delta t/\tau_{\mathrm{min}} = 3$, and recovers the correct scaling of the relaxation times for various time interval to relaxation time ratios (Fig. S18b).

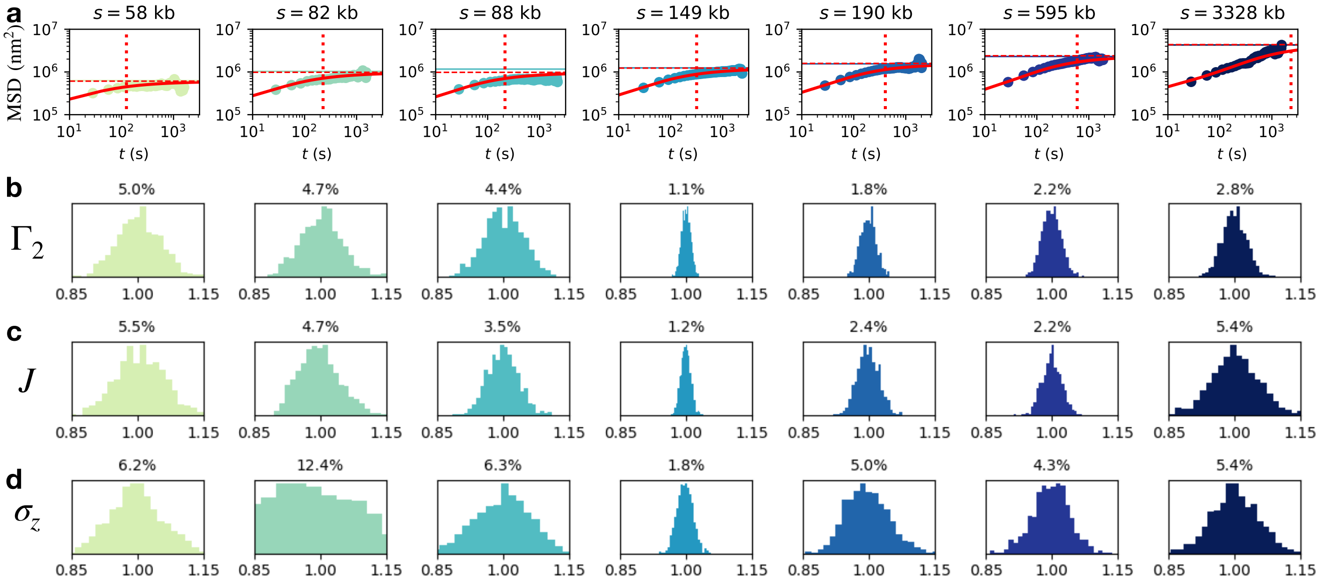

Figure S17: **Two-locus MSD fitting and parameter inference.** **a**, Two-locus MSDs $M_2(t)$ for each genomic separation. Solid lines in colors show the value of $2\langle R^2\rangle$. Solid red lines show the fitted $M_2$ with measurement error correction. Dashed red lines show the fitted asymptote $2J$ and dotted red lines show inferred relaxation time $\tau$. **b-d**, Distributions of the bootstrapped estimates for each parameter, normalised by the maximum a posterior estimate of each parameter, $\theta/\theta_{\mathrm{MAP}}$.

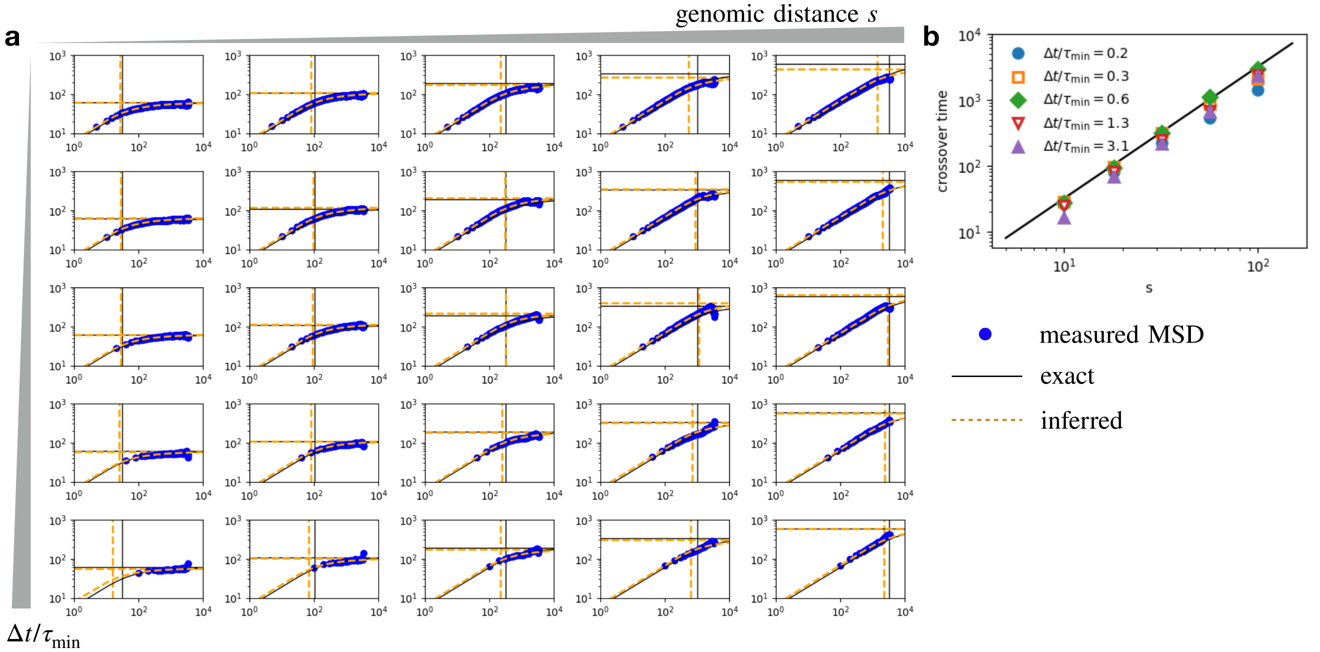

Figure S18: **Testing the relaxation time inference on simulated ideal Rouse chains.** **a**, Two-locus MSDs $M_2(t)$. Blue symbols: measured two-locus MSD from the simulations. Black solid lines: exact values of the MSD, the relaxation time, and the plateau $2\langle R^2\rangle$. Orange dashed lines: inferred values of these three quantities. Horizontal lines show the value of $2\langle R^2\rangle$, vertical lines show inferred relaxation time $\tau$. Rows correspond to the five tested $\Delta t/\tau_{\mathrm{min}}$-ratios listed in the legend of panel b. **b**, Exact (line) and inferred (symbols) relaxation time scaling for varying $\Delta t/\tau_{\mathrm{min}}$.

An alternative, simplified way to obtain an estimate of the relaxation time is by fitting the early time two-locus MSD and measuring the intercept with the measured value of $2\langle R^2\rangle$ (Fig. S19a). This intercept is then given by $\tau \sim (\langle R^2\rangle/\Gamma_2)^{1/\beta}$. We test this simplified approach both with $\beta \approx 0.5$ as measured

| $s$ (kb) | $N_{\text{embryos}}$ | $N_{\text{nuclei}}$ | $\hat{\sigma}_z$ (nm) | $J$ (nm$^2$) | $\Gamma_2$ (nm$^2$s$^{-1/2}$) |
|---|---|---|---|---|---|
| 58 | 7 | 55 | $200 \pm 13$ | $(2.6 \pm 0.1) \times 10^5$ | $(2.4 \pm 0.1) \times 10^4$ |
| 82 | 4 | 59 | $177 \pm 10$ | $(4.5 \pm 0.2) \times 10^5$ | $(3.0 \pm 0.1) \times 10^4$ |
| 88 | 5 | 58 | $210 \pm 26$ | $(4.6 \pm 0.2) \times 10^5$ | $(3.1 \pm 0.1) \times 10^4$ |
| 149 | 30 | 579 | $204 \pm 4$ | $(5.73 \pm 0.07) \times 10^5$ | $(3.23 \pm 0.04) \times 10^4$ |
| 190 | 16 | 166 | $220 \pm 10$ | $(7.4 \pm 0.2) \times 10^5$ | $(3.70 \pm 0.07) \times 10^4$ |
| 595 | 16 | 164 | $213 \pm 9$ | $(1.1 \pm 0.03) \times 10^6$ | $(4.7 \pm 0.1) \times 10^4$ |
| 3328 | 10 | 179 | $340 \pm 10$ | $(2.1 \pm 0.1) \times 10^6$ | $(4.35 \pm 0.09) \times 10^4$ |

Table S5: **Statistics of the trajectory data sets of all constructs**. From left to right, the columns denote: (i) The number of tracked embryos. (ii) The total number of tracked nuclei. (iii) The estimated measurement error in the axial direction, obtained from the MSD fitting. (iv) The fitted value of $J$. (v) The fitted value of $\Gamma_2$.

from the single-locus MSD. We find that this simplified approach recovers the strongly reduced scaling exponent compared to the ideal Rouse expectation of $\gamma = 2$ (Fig. S19b).

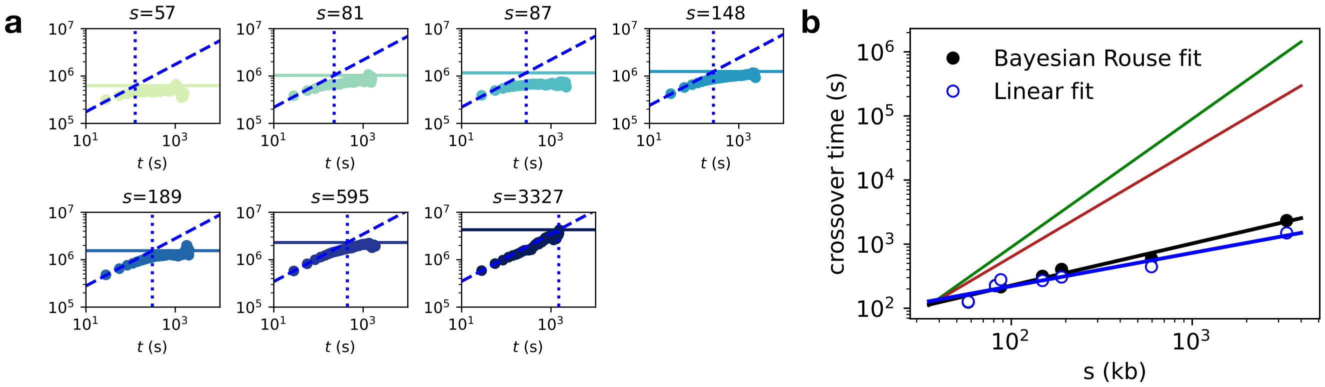

Figure S19: **Relaxation time estimation with simple intercept fitting.** **a**, Two-locus MSDs $M_2(t)$ for each genomic separation. Solid lines show the value of $2\langle R^2\rangle$. Dashed lines are fits of $2\Gamma_2 t^{0.5}$. Dotted lines show the intercept of the fit with $2\langle R^2\rangle$, providing an estimate for the relaxation time $\tau$ for each of the two scenarios. **b**, Estimated cross-over times from both the Bayesian inference of the full Rouse dependence used in the main text (Eq. (S28), black filled dots) and the linear log-fit (blue open symbols).

#### 4.2 Two-locus autocorrelation collapse

We make use of the predicted collapse of the two-locus auto-correlation (ACF) (Eq. (S57)) to test the quality of the collapse for the relaxation time exponents for various polymer models and compare them to the anomalous scaling collapse S20. As expected, we find that the correlations do not collapse for the ideal and crumpled chain relaxation time scaling exponents, but they do exhibit an approximate collapse when using the measured relaxation time scaling $\gamma = 0.7$.

#### 4.3 Velocity cross-correlations

**Error-corrected correlation estimators**: To measure the correlations in the joint motion of two loci in absolute motion (within the frame of reference of the cell nucleus), we use the cross-correlations of

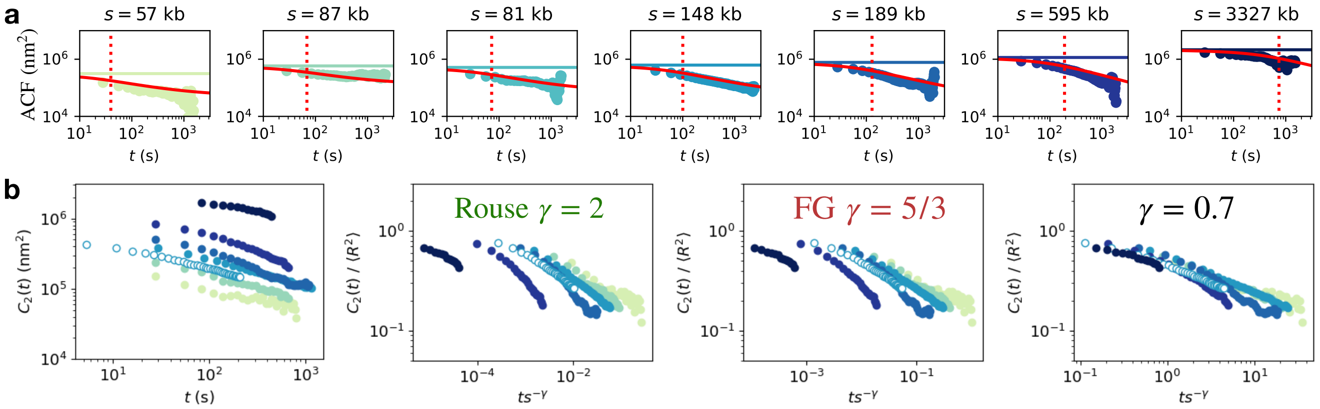

Figure S20: **Two-locus autocorrelation collapse.** **a**, Two-locus ACFs $C_2(t)$ for each genomic separation. Solid lines in color: plateau value $2\langle R^2\rangle$. Red line: ACF expected based on the fitted MSDs, $C_2(t) = (2\langle R^2\rangle - M_2(t))/2$. **b**, All ACFs plotted in absolute units without collapse. **c**, ACFs collapsed by the measured value of $\langle R^2\rangle$ along the y-axis and with the ideal chain relaxation time scaling $\gamma = 2$. **d**, Collapse using the crumpled chain relaxation time scaling $\gamma = 5/3$. **e**, Collapse using the measured anomalous relaxation time scaling $\gamma = 0.7$.

velocities, defined by

$$C_{vv}^{(\delta)}(t) = \langle \mathbf{v}_i^{(\delta)}(t_0) \cdot \mathbf{v}_j^{(\delta)}(t_0 + t)\rangle_{t_0} \tag{S29}$$

These correlations are challenging to measure as the absolute motion is affected by global motion of the cell nuclei. Since the syncitial nuclei in the early *Drosophila* embryo are relatively stable and only exhibit slow affine translations, we correct for global motion in the $xy$-direction and estimate velocity cross-correlations for the $x$ and $y$ components as done for single locus MSDs (section 3). To further correct for the random localization errors in the nucleus tracking, we derive error-corrected correlation estimators. As we show in Fig. S3d, measurement errors in the nucleus tracking are uncorrelated in time. Importantly, however, even time-uncorrelated measurement errors can impact the estimated velocity cross-correlations, because the nucleus movement is subtracted from the motion of both loci, which may lead to spurious additional correlations in their apparent velocities. To overcome this issue, we derive the noise correction terms of the velocity cross-correlations, providing correlation estimators that are unbiased with respect to measurement noise.

Specifically, throughout, we consider the position of locus $i$ in the frame of reference in the nucleus,

$$x_i = r_i - r_n \tag{S30}$$

where $r_i$ is the locus position in the lab frame and $r_n$ is the nucleus position in the lab frame. Our measurement of these positions contains measurement noise, and thus

$$\hat{r}_i = r_i + \sigma_r \eta_i(t) \tag{S31}$$

$$\hat{r}_n = r_n + \sigma_n \xi(t) \tag{S32}$$

where we assume temporally uncorrelated measurement errors, i.e. $\langle \eta_i(t)\rangle = \langle \xi(t)\rangle = 0$ and $\langle \eta_i(t)\eta_j(t')\rangle = \delta_{ij}\delta(t-t')$ and $\langle \xi(t)\xi(t')\rangle = \delta(t-t')$. Here $\sigma_r$ is the measurement noise in locus tracking and $\sigma_n$ is the measurement error i the nucleus tracking. Combining these assumptions, we find

$$\hat{r}_i = r_i + \sigma_r \eta_i(t) - \sigma_n \xi(t) \tag{S33}$$

and thus

$$\hat{v}_i^{(\delta)}(t) = (\hat{x}_i(t+\delta) - \hat{x}_i(t))/\delta \tag{S34}$$

$$= v_i^{(\delta)}(t) + \frac{\sigma_r}{\delta}[\eta_i(t+\delta) - \eta_i(t)] + \frac{\sigma_n}{\delta}[\xi(t+\delta) - \xi(t)] \tag{S35}$$

Multiplying out, we can therefore find the correction terms to various correlation functions. First, we consider the correlation of the locus velocity with the nucleus velocity:

$$\langle \hat{v}_i^{(\delta)}(t)\hat{v}_n^{(\delta)}(0)\rangle = \langle v_i^{(\delta)}(t)v_n^{(\delta)}(0)\rangle + \begin{cases} -2\sigma_n^2/\delta^2 & t=0 \\ \sigma_n^2/\delta^2 & t=\delta \\ 0 & \text{otherwise} \end{cases} \tag{S36}$$

This shows that under the assumption of uncorrelated noise this correlation should vanish except for $\tau = \{0, \delta\}$, which we verify on the experimental data, validating the assumption of time-uncorrelated tracking errors (Fig. S3d).

Similarly, in the velocity cross-correlations, most terms due to measurement noise average to zero, but the terms due to nucleus tracking errors do not, as the same random error affects both loci:

$$\hat{C}_{vv}(t) = \langle \hat{v}_i^{(\delta)}(t)\hat{v}_j^{(\delta)}(0)\rangle \tag{S37}$$

$$= C_{vv}(t) + \begin{cases} 2\sigma_n^2/\delta^2 & t=0 \\ -\sigma_n^2/\delta^2 & t=\delta \\ 0 & \text{otherwise} \end{cases} \tag{S38}$$

Similarly, for the velocity autocorrelation, we obtain

$$\hat{C}_{v}(t) = \langle \hat{v}_i^{(\delta)}(t)\hat{v}_i^{(\delta)}(0)\rangle \tag{S39}$$

$$= C_{v}(t) + \begin{cases} 2(\sigma_n^2+\sigma_r^2)/\delta^2 & t=0 \\ -(\sigma_n^2+\sigma_r^2)/\delta^2 & t=\delta \\ 0 & \text{otherwise} \end{cases} \tag{S40}$$

Note that the above derivation was done in 1D, for 3D all contributions simply add up, meaning that $\sigma_n^2 \to \sigma_{n,x}^2 + \sigma_{n,y}^2 + \sigma_{n,z}^2$. Importantly, these corrections show that errors scale with the time-interval $\delta$ at which velocities are calculated. Consequently, we find that the estimated velocity cross-correlations are more strongly affected at small $\delta$. To identify how large $\delta$ should be for our estimate to be robust against measurement noise, we plot the cross-correlation intercepts $C_{vv}(0)/C_v(0)$ as a function of $\delta$ for different values of the estimated measurement error. Based on the fitted measurement errors from the two-locus MSDs (section 4.1), we expect $\sigma \approx 200\mu\text{m}$. We find that for $\delta$ in the range of 5 to 60 seconds, the estimated cross-correlations are strongly affected by realistic measurement error amplitudes (Fig. S21a). In contrast for $\delta > 60$ s, the correlations are largely insensitive to realistic measurement errors for all genomic separations (Fig. S21b). The larger $\delta$, however, the smaller the sample size of available time-windows in a finite trajectory, resulting in larger random errors. As a tradeoff, we therefore focus our analysis of velocity cross-correlations on $\delta = 90$ s, but additionally show variation with $\delta$ in Fig. S22.

**Theory prediction of velocity cross-correlations**: The velocity cross-correlation of two loci in the ideal chain Rouse model have been calculated analytically in ref. [19]. Importantly, in the limit $s \ll N$, where $N$ is the total polymer length, and $\delta \ll \tau_N$, where $\tau_N$ is the relaxation time of the whole polymer, the cross-correlations of two loci separated by genomic separation $s$ are completely determined by the dimensionless ratio $\delta/\tau$, where $\tau$ is the relaxation time of the segment $s$. The correlations are calculated using the tabulated values in ref. [19] (using viscoelastic exponent $\alpha = 1$).

Thus, we can predict the velocity cross-correlations expected based on the inferred relaxation times without any free fitting parameters. We predict the correlations for varying $s$ based on the scaling of

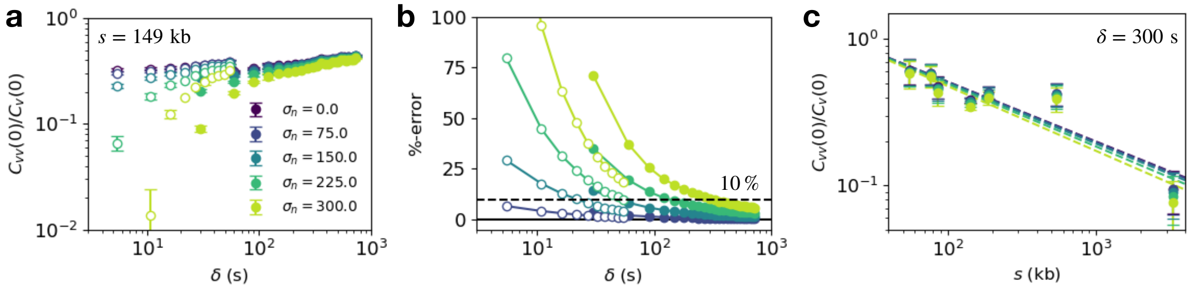

Figure S21: **Error-correction of the velocity cross-correlations.** **a**, Normalized velocity cross-correlations as a function of the time-interval $\delta$ for $s = 149$ kb. Open symbols: data acquired at imaging rate $\Delta t = 5.4$ s. Closed symbols: data acquired at imaging rate $\Delta t = 30$ s. Different colors correspond to error-corrected correlations assuming different measurement errors (measured in µm), were we take $\sigma_n = \sigma_r = \sigma$. We plot the error-free correlations $C_{vv}(0)/C_v(0)$ calculated from $\hat{C}_{vv}(0)/\hat{C}_v(0)$ according to Eqs. (S37), (S39). **b**, Fractional error in the velocity correlations in panel a due to the measurement error contribution, in percent. The dashed line indicates the 10% mark. For $\delta > 90$s, the errors are $< 10\%$ for all values of $\sigma$. Colors correspond to the legend in panel a. **c**, Normalized velocity cross-correlations as a function of the genomic separation $s$ for $\delta = 90$ s. Colors correspond to the legend in panel a.

relaxation times $\tau = As^{\gamma}$. First, we predict the expected correlations assuming an ideal chain Rouse scaling of the relaxation time, i.e. $\gamma = 2$. In main text Fig. 4h, we use the prefactor $A$ based on the relaxation time of the shortest genomic separation to the largest genomic separation, i.e. $A^{(s)}_{\text{ideal}} = \tau/s^2$. Here, we additionally demonstrate the ideal chain prediction for varying $A$, showing that the prediction fails for any possible value of $A$ (Fig. S22a).

To predict the correlations expected based on the anomalous scaling that we find in experiments, we use the measured value of $A = 1.89$ s kb$^{-\gamma}$ and $\gamma = 0.7$. Thus, both parameters are fixed based on the experimental data with no additional fitting, giving the blue line in Fig. S22a. Based on the arguments in the previous section, suggesting that larger time-intervals $\delta$ are less affected by measurement errors, we use $\delta = 300$ s in the main text figure. Here, we additionally show the correlations for varying values of $\delta$ (Fig. S22b,d). The functional dependence of the correlations on $\delta$ is shown in Fig. S22c. We find that the correlations are generally well predicted by the theory curves that are completely constrained by the inferred relaxation time, with no further fitting involved.

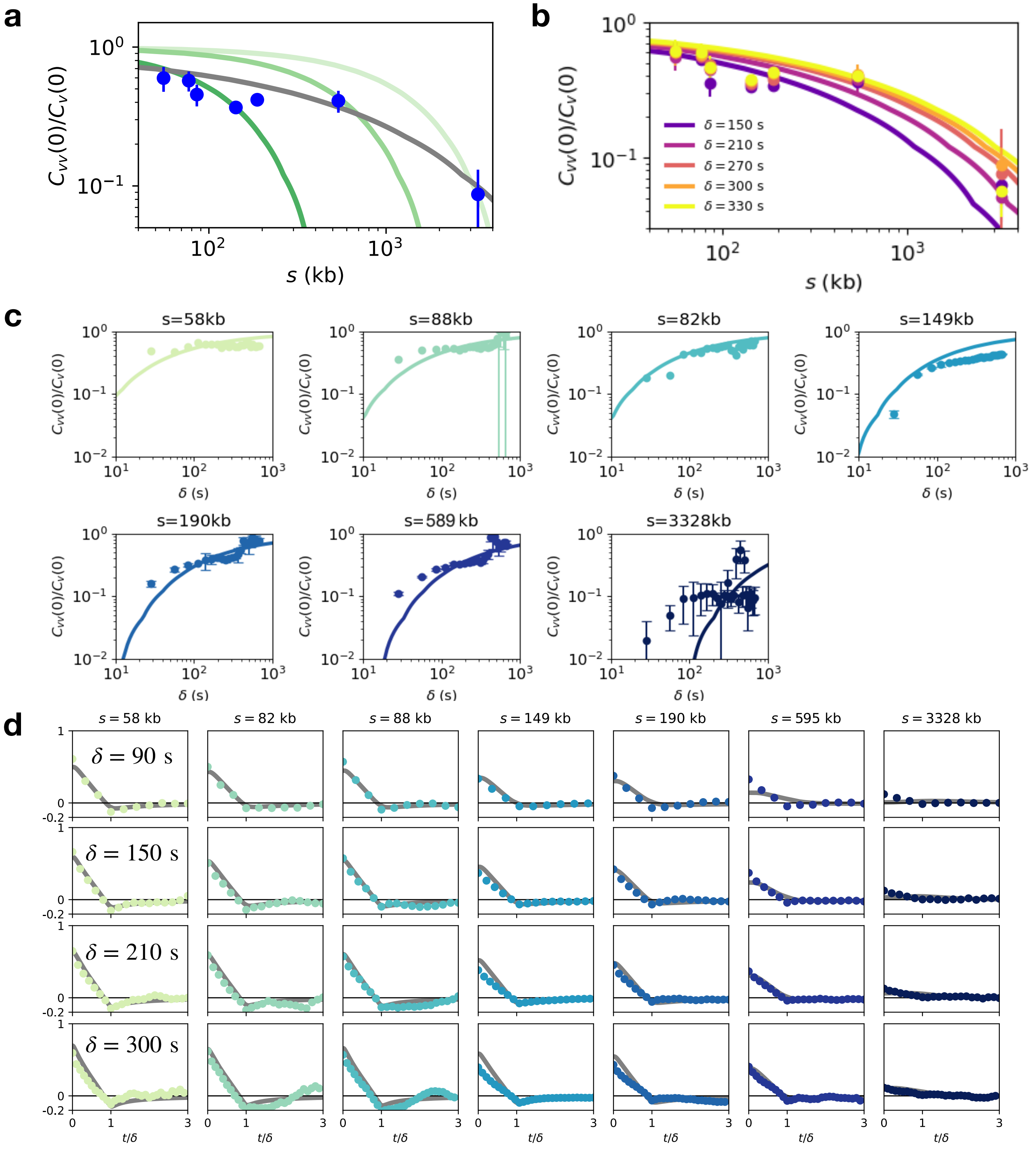

Figure S22: **Theory prediction of velocity cross-correlations.** **a**, Comparison of experimental correlations to predictions based on the ideal Rouse scaling $\tau = As^2$, for varying values of $A$, calculated to match increasing values of $s = \{58\text{kb}, 595\text{kb}, 3.3\text{Mb}\}$, using $A^{(s)}_{\text{ideal}} = \tau/s^2 = \{0.023, 8.1 \times 10^{-4}, 1.01 \times 10^{-4}\}$ s kb$^{-2}$ (green curves). **b**, Anomalous scaling prediction compared to experimental data for various time-intervals $\delta$. **c**, Velocity cross-correlations as a function of the time-interval $\delta$ (colors), as predicted by the anomalous relaxation time scaling (solid lines) compared to experiments (symbols) **d**, Velocity cross-correlation functions $C_{vv}(t)/C_v(0)$ as a function of $t/\delta$ for varying $\delta$ for each genomic separation. All errorbars correspond to bootstrap errors.

### 5 Two-locus dynamics: theory

In this section, we provide a synthesis of the literature on the scaling exponents measured in this paper, and explain the theoretical arguments underlying our analysis and interpretation two-locus dynamics.

**Two-locus MSD:** Here, we provide details of how we quantify the joint dynamics of two coupled DNA loci using the two-locus MSD:

$$M_2(t) = \left\langle \left|\mathbf{R}_{ij}(t_0+t) - \mathbf{R}_{ij}(t_0)\right|^2 \right\rangle_{t_0} \tag{S41}$$

i.e. the MSD of the 3D interlocus distance vector $\mathbf{R}_{ij}(t) = \mathbf{r}_1(t) - \mathbf{r}_2(t)$, where $\mathbf{r}_{1,2}$ are the 3D positions of the two loci.

The two-locus MSD gives insight into the coupled dynamics of the two loci, with two simple limits: at small times $t \ll \tau$, we expect the two-locus MSD to simply reflect the independent diffusion of the two loci, while at large times $t \gg \tau$, the MSD approaches a plateau given by the equilibrium size of the chain. Here, $\tau$ is the relaxation time which quantifies the cross-over between the two regimes.

In the large-time regime, the chain segment connecting the two loci has fully relaxed (under the assumptions that the system reaches a steady state), meaning that the two-locus MSD is given by the equilibrated size of the chain

$$M_2(t \to \infty) \to 2\langle R^2 \rangle \tag{S42}$$

where $\langle R^2 \rangle = \langle |\mathbf{R}_{ij}|^2 \rangle$. For a fractal polymer chain configuration, the average 3D RMS-distance $\sqrt{\langle R^2 \rangle}$ scales with the genomic separation $s$:

$$\sqrt{\langle R^2 \rangle} \sim s^{\nu} \tag{S43}$$

where the *static* exponent $\nu = 1/d$ is determined by the fractal dimension $d$ of the polymer. The fractal dimension is $d = 2$ for an ideal chain polymer, and $d = 3$ for a crumpled chain.

In the small-time regime, the chain segment connecting the two loci has not had sufficient time to relax, and the two loci therefore move independently, meaning that the two-locus MSD is given by the sum of the single-locus MSDs:

$$M_2(t \ll \tau) \approx 2M_1(t) = 2\Gamma t^{\beta} \tag{S44}$$

Importantly, the dynamic exponent $\beta$ can be related to the static exponent $\nu$, and thus the fractal dimension $d$ under specific assumptions for the dynamics of the polymer, which we detail below. To avoid confusion, we try to be specific about which assumptions relate to the statics and which to the dynamics. For statics, we use the terms "ideal" and "crumpled chain statistics" for fractal dimensions $d = 2$ and $d = 3$, respectively. For the dynamics of such ideal or crumpled chains, we use the term "generalized Rouse assumption" to refer to the assumption of independent viscous friction on each monomer, consistent with the terminology in [22].

**Generalized Rouse approach:** Within the framework of a generalized Rouse approach, $\beta$ can be related to the static properties for arbitrary fractal dimension $d$ [22–24]:

$$\beta = \frac{2}{2+d} \tag{S45}$$

This is derived as follows: at a time-lag $t$, due to stress propagation, a region of spatial size $\ell \sim t^{\beta/2}$ behaves effectively as a single monomer. Then, the key assumption is the so-called 'free-draining' limit, i.e.

only local friction on every monomer and no hydrodynamic interactions. Thus, the effective diffusion coefficient $D$ of this region then scales with the number of monomers contained within the region, which is determined by the fractal dimension: $D \sim \ell^{-d}$. Therefore, the MSD of the region is $Dt \sim t^{1-\beta d/2}$, which yields Eq. (S45). Eq. (S45) then yields $\beta = 1/2$ for $d = 2$ and $\beta = 2/5$ for the crumpled chain ($d = 3$).

**Zimm model:** The Zimm model describes polymers that have a hydrodynamic self-interaction [11], meaning that the monomer beads do not only interact with their nearest neighbours through the springs linking them, but through long-ranged hydrodynamic interactions. Within the Zimm model, the dynamic exponent is independent of the fractal packing dimension $d$ and is given by $\beta = 2/3$.

**Relaxation time scaling:** To quantify the time-scale on which the cross-over between the two regimes (Eq. (S42) and Eq. (S44)) occurs, we focus on the relaxation time $\tau$. The relaxation time can be estimated as the time taken for the two loci to diffuse their typical distance of separation [22]. In terms of the two-locus MSD, this time-scale is therefore given by the intersection of the diffusive part (Eq. (S44)) and the long-time plateau (Eq. (S42)):

$$2\Gamma_2 \tau^{\beta} \sim 2\langle R^2 \rangle \tag{S46}$$

Based on this relationship, we can infer the relaxation times from the experimental two-locus MSD curves, as shown in detail in section 4.1. Furthermore, Eq. (S46) predicts the scaling of the relaxation time with genomic separation for different fractal dimensions for both the generalized Rouse and the Zimm models:

$$\tau^{\beta} \sim s^{2/d} \quad \rightarrow \quad \tau \sim s^{\gamma} \text{ with } \gamma = 2/(\beta d) \tag{S47}$$

We therefore obtain the classical relaxation time scaling for the ideal chain Rouse model:

$$\tau^{1/2} \sim s \quad \rightarrow \quad \tau \sim s^2 \tag{S48}$$

For a crumpled chain described by the generalized Rouse model, both the diffusion exponent and the packing exponent change, and we therefore obtain

$$\tau^{2/5} \sim s^{2/3} \quad \rightarrow \quad \tau \sim s^{5/3} \tag{S49}$$

Thus, the relaxation time exhibits a weaker scaling with genomic separation in the crumpled chain model (exponent $\approx 1.66$) than in the ideal chain model (exponent 2). These two exponents are shown as the theory predictions in main text Fig. 4.

We can make the same arguments based on the exponents of the Zimm model for ideal and crumpled chains. Thus, for the ideal chain Zimm model,

$$\tau^{2/3} \sim s \quad \rightarrow \quad \tau \sim s^{3/2} \tag{S50}$$

For a crumpled chain described by the Zimm model, we obtain

$$\tau^{2/3} \sim s^{2/3} \quad \rightarrow \quad \tau \sim s^1 \tag{S51}$$

These various results are summarised in Table S6.

| statics \ dynamics | Rouse | Zimm |
|---|---|---|
| general $d$ | $\beta = 2/(2+d)$<br>$\gamma = (2+d)/d$ | $\beta = 2/3$<br>$\gamma = 3/d$ |
| ideal chain<br>$d = 2$ | $\beta = 1/2$<br>$\gamma = 2$ | $\beta = 2/3$<br>$\gamma = 3/2$ |
| crumpled chain<br>$d = 3$ | $\beta = 4/5$<br>$\gamma = 5/3$ | $\beta = 2/3$<br>$\gamma = 1$ |

Table S6: **Overview of exponents $\beta, \gamma$ expected for various combinations of statics (ideal/crumpled) and dynamics (Rouse/Zimm).**

**Two-locus autocorrelation function:** Besides the two-locus MSD, a second quantity that quantifies the joint dynamics of two loci based on the interlocus distance vector is the two-locus autocorrelation, given by

$$C_2(t) = \left\langle \mathbf{R}_{ij}(t_0 + t) \cdot \mathbf{R}_{ij}(t_0) \right\rangle_{t_0} \tag{S52}$$

The quantities $M_2$ and $C_2$ contain the same information, since they are directly related to each other through the relationship [14]

$$M_2(t) = 2(C_2(0) - C_2(t)) \tag{S53}$$

However, the function $C_2$ gives access to the scaling exponents that characterize the approach to equilibration in the long time limit $t \gg \tau$, which are 'washed out' in the cross-over to the plateau in $M_2$.

Specifically, for $C_2$, the short time behaviour ($t \ll \tau$) is determined by the chain size:

$$C_2(t \to 0) = \langle R^2 \rangle \tag{S54}$$

The asymptotic power-law behaviour of the two-locus correlation function in the limit $t \ll \tau$ has been derived based within the generalized Rouse framework [22], predicting that it decays with a power-law whose exponents depend on the fractal dimension:

$$C_2(t) \sim t^{-\lambda} \tag{S55}$$

where

$$\lambda = 2\frac{d-1}{2+d} \tag{S56}$$

which gives $\lambda = 1/2$ for ideal and $\lambda = 4/5$ for crumpled chains. Note that this derivation additionally assumes $s \ll N$, where $N$ is the total length of the chromosome. In our system (*Drosophila* chromosome 2L), $N = 25$ Mb, significantly larger than the typical genomic separations we probe (58 kb – 3.3 Mb). Furthermore, we do not see any signatures of whole chain relaxation in the single-locus MSDs (Fig. S14), suggesting this assumption is valid. Thus, the two-locus autocorrelation can be used to quantify the approach to equilibration of the system, which is separate to the small-time scaling of the two-locus MSD, quantified by exponent $\beta$, which is determined by the independent diffusion of the loci.

Finally, we can use the two-locus autocorrelation as an additional way to infer the relaxation times by collapsing the autocorrelations onto a single master curve. Specifically, we expect the correlations to collapse onto the mastercurve [22]

$$\mathcal{C}_2(\tilde{t}) \sim s^{-2/d} C_2(ts^{-\gamma}) \tag{S57}$$

where $\gamma$ is the exponent of the relaxation time, $\tau \sim s^{\gamma}$. Thus, by quantifying the collapse of $C_2$ for varying $s$, we can identify the scaling exponent $\gamma$ (see section 4.2).

### 6 Comparison to mESC data

To compare our quantitative results to previous measurements of live imaging of pairs of DNA loci, we make a side-by-side comparison of our data set with that in ref. [14], where CTCF sites in the *Fbn2* locus imaged in mouse embryonic stem cells (mESCs) were tagged with a genomic separation of 515 kb between fluorescent tags (C36 line). We compare our closes genomic separation to this data (595 kb), and find that both average distances and diffusive time-scales are much higher in our system. Specifically, while the average distance for mESCs is $\langle R \rangle \approx 0.35\mu m$, we find $\langle R \rangle \approx 1.05\mu m$, a factor 3 larger. Furthermore, the two-locus diffusivity $\Gamma_2 = 0.0024 \mu m^2 s^{-1/2}$ for mESCs, and $\Gamma_2 = 0.045 \mu m^2 s^{-1/2}$ in our system, almost 20-fold larger. As a result, despite the larger physical distance between the two loci, the relaxation time is almost an order of magnitude shorter than observed in ref. [14].

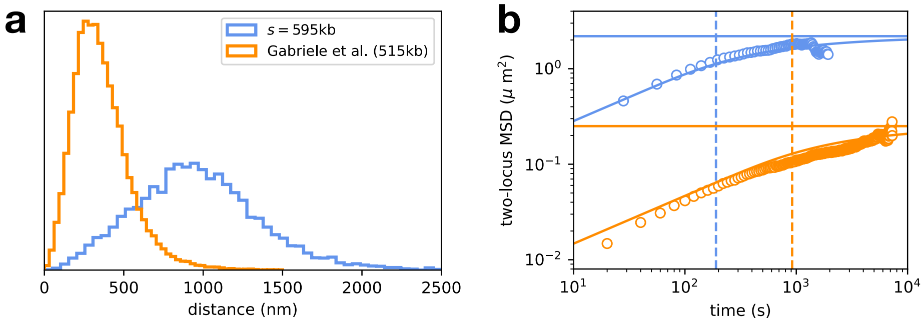

Figure S23: **Two-locus analysis compared to data from ref. [14].** **a**, Probability distributions of the inter-locus distances $R_{ij}$. Blue: our data for $s$ = 595 kb (all states), orange: data of CTCF sites in the *Fbn2* locus imaged in mouse embryonic stem cells (mESCs), with a genomic separation of 515 kb (C36 wildtype line) [14]. **b**, Localization-error corrected two-locus MSDs for both data sets (open symbols). Solid line: Fit of Eqn. (S28). Solid horizontal line: $2\langle R^2 \rangle$. Dashed vertical line: fitted value of the relaxation time $\tau$.

### 7 Data and materials availability

Code for this paper is available at `https://github.com/dbrueckner/DynamicChromosome`. The raw trajectory data is available at: `https://doi.org/10.5281/zenodo.7616203`. Additional code and software used in this paper are listed in Table S7.

| Name | Reference | URL |
|---|---|---|
| numpy 1.22.0 | [25] | `https://numpy.org/` |
| scipy 1.4.1 | [26] | `https://scipy.org/` |
| matplotlib 3.1.3 | [27] | `https://matplotlib.org/` |
| tracklib | [14] | `https://github.com/SGrosse-Holz/tracklib` |
| Matlab 2020b | – | `https://mathworks.com/products/matlab.html` |

Table S7: **List of software used**.

### 8 Movie description

**Supplementary Movies S1-S7**
Three-color live imaging of two loci on the second Drosophila chromosome separated by increasing genomic distances of $\{57.8, 81.7, 87.5, 148.7, 189.6, 595.1, 3327.7\}$ kb. The embryo is 2 h old. Movie starts from 27 min in cell cycle 14 and ends at 55 min. Blue spots mark eve-MS2 transcription as well as the physical position of the eve locus and eve enhancers. Green spots are from the parS sequence, which marks the location of the reporter construct. Red spots mark transcriptional activity from the PP7 reporter. Scale bar: 5 µm.

[26] P. Virtanen, R. Gommers, T. E. Oliphant, M. Haberland, T. Reddy, D. Cournapeau, E. Burovski, P. Peterson, W. Weckesser, J. Bright, S. J. van der Walt, M. Brett, J. Wilson, K. J. Millman, N. Mayorov, A. R. J. Nelson, E. Jones, R. Kern, E. Larson, C. J. Carey, İ. Polat, Y. Feng, E. W. Moore, J. VanderPlas, D. Laxalde, J. Perktold, R. Cimrman, I. Henriksen, E. A. Quintero, C. R. Harris, A. M. Archibald, A. H. Ribeiro, F. Pedregosa, P. van Mulbregt, A. Vijaykumar, A. P. Bardelli, A. Rothberg, A. Hilboll, A. Kloeckner, A. Scopatz, A. Lee, A. Rokem, C. N. Woods, C. Fulton, C. Masson, C. Häggström, C. Fitzgerald, D. A. Nicholson, D. R. Hagen, D. V. Pasechnik, E. Olivetti, E. Martin, E. Wieser, F. Silva, F. Lenders, F. Wilhelm, G. Young, G. A. Price, G.-L. Ingold, G. E. Allen, G. R. Lee, H. Audren, I. Probst, J. P. Dietrich, J. Silterra, J. T. Webber, J. Slavič, J. Nothman, J. Buchner, J. Kulick, J. L. Schönberger, J. V. de Miranda Cardoso, J. Reimer, J. Harrington, J. L. C. Rodríguez, J. Nunez-Iglesias, J. Kuczynski, K. Tritz, M. Thoma, M. Newville, M. Kümmerer, M. Bolingbroke, M. Tartre, M. Pak, N. J. Smith, N. Nowaczyk, N. Shebanov, O. Pavlyk, P. A. Brodtkorb, P. Lee, R. T. McGibbon, R. Feldbauer, S. Lewis, S. Tygier, S. Sievert, S. Vigna, S. Peterson, S. More, T. Pudlik, T. Oshima, T. J. Pingel, T. P. Robitaille, T. Spura, T. R. Jones, T. Cera, T. Leslie, T. Zito, T. Krauss, U. Upadhyay, Y. O. Halchenko, Y. Vázquez-Baeza, and S. . Contributors, "SciPy 1.0: Fundamental algorithms for scientific computing in Python," *Nature Methods*, vol. 17, no. 3, pp. 261–272, 2020.

[27] J. D. Hunter, “Matplotlib: A 2D Graphics Environment,” *Computing in Science & Engineering*, vol. 9, no. 3, pp. 90–95, 2007.
